# Interactions of Upstream and Downstream Promoter Regions with RNA Polymerase are Energetically Coupled and a Target of Regulation in Transcription Initiation

**DOI:** 10.1101/2020.05.13.070375

**Authors:** Robert P. Sosa, Alfredo J. Florez-Ariza, César Díaz-Celis, Bibiana Onoa, Alexandre Cassago, Paulo S. L. de Oliveira, Rodrigo V. Portugal, Daniel G. Guerra, Carlos J. Bustamante

## Abstract

During transcription initiation, the RNA polymerase holoenzyme (RNAP) and the promoter form an open complex. For many promoters, this interaction involves upstream DNA wrapping, downstream promoter bending, and associated large-scale protein rearrangements. Although these processes have been reported across the life kingdom, their structure, energetics, and role in transcription remain an area of active research. Using optical tweezers, we find that these processes become energetically and reversibly coupled after the formation of the open promoter complex, providing the main contribution to their stability. Using electron microscopy and single particle analysis, we find that the interaction encompasses from positions −76 to +18 along the template, that it involves an overall DNA bent angle of ~245°, and that the upstream wrapping is enabled by interactions between the C-terminal domains of RNAP’s alpha subunits and proximal and middle upstream promoter regions. The energy associated with upstream wrapping, downstream bending and its coupling to downstream rearrangements does not require specific upstream promoter sequence, and correlate positively with the rate of transcription DNA bubble formation as reported by a real-time fluorescence assay. Our results suggest that the coupling between upstream and downstream events are part of a *cis*-regulatory network established after the opening of the DNA bubble, that could furnish a control mechanism of gene expression by protein factors and regulatory metabolites.

**Summary:** The first step of gene expression involves transcription of DNA into RNA by RNA polymerase (RNAP). RNAP recognizes a promoter sequence forming the transcriptionally active open complex. For several promoters, DNA wraps around the RNAP. We find that upstream wrapping contacts are energetically coupled and occur cooperatively with downstream rearrangements in the open complexes, providing the largest contribution to their stability. We also determined that upstream wrapping is enabled by interactions between non-specific upstream promoter regions and RNAP α subunit C-terminal domains. Significantly, the strength of these contacts correlates with the rate of DNA bubble opening, and is regulated by factors such as the transcriptional regulator ppGpp. We suggest that any modulator altering upstream wrapping and downstream rearrangements could finely tune gene expression in response to the needs of the cell

## Introduction

Transcription in prokaryotes is initiated when the RNA polymerase holoenzyme (RNAP) recognizes specialized DNA promoter sequences. Upon enzyme binding, the DNA-protein complex undergoes a series of conformational changes—upstream wrapping, downstream bending, accommodation of downstream DNA within the cleft formed by the β′-jaw/β’-clamp domains of RNAP, transcription DNA bubble formation, folding of downstream mobile elements (DME) in the β′-jaw, and the repositioning of the σ^70^_1.1_ domain—leading to the formation and stabilization of the open complex (RPo) [1], [2]. Promoter sequences and several transcription regulators (proteins and metabolites such as guanosine tetra-phosphate, ppGpp) [3] combine to produce a context-dependent kinetic pathway for open complex formation, where the relative stabilities of the various intermediates in this process are specific of each promoter [1], [2]. Before the enzyme can escape from the promoter into processive elongation, the open complex undergoes multiple cycles of abortive initiation, releasing short RNA products. Accordingly, open complex formation and its escape from the promoter constitute major regulatory checkpoints in transcription throughput.

Most open complex studies have focused on the core promoter elements (such as −10 and −35 boxes and their direct vicinity), which are involved in sequence-dependent recognition by the enzyme, as well as in DNA bubble formation [1], [2], [4]–[6]. However, various lines of evidence indicate that a more extended portion of the promoter DNA wraps around the enzyme, both in prokaryotic and in mammalian pre-initiation open complexes [7], [8]. Specifically, fluorescence (λ*p_R_*) [9] and footprinting assays (λ*p_R_*, *lacUV*5, *gal*P1, T7A1, *mal*T, Bla, *Alu*156) [8], [10]–[15], electron microscopy (EM) and atomic-force microscopy (AFM) (adenovirus major late promoter, λ*p_R_*, *rrnB* P1, tyrT, *hdeAB*p,) [7], [16]–[19], and crosslinking (adenovirus major late promoter) [20], indicate that 50, 100 or more base pairs upstream of the transcription start site wrap around the enzyme. AFM data showed that wrapping of the λ*p_R_* promoter may involve interactions of the alpha carboxy terminal domains (αCTD) of the RNAP with upstream AT-rich sequences [21]. Although upstream wrapping *per se* contributes little to open complex stability [22], [23], these DNA contacts with the enzyme are thought to play a regulatory role in DNA bubble formation and downstream promoter-RNAP interactions in open complexes [22], [24]–[27]. Also large-scale rearrangements associated to downstream bending, such as folding of downstream mobile elements in the β′-jaw (DME), and the repositioning of the σ^70^_1.1_ domain, have been proposed to stabilize the open complex [1], [6], [28]. Thus, upstream wrapping and downstream rearrangements may play an important role in the formation and stabilization of open complexes which can, in turn, be subjected to modulation by metabolites such as ppGpp or transcription factors such as DksA [3]. Despite the critical importance of these extended interactions, no direct measurement of their strength has been explored in open complexes.

Here, we use optical tweezers to study the strength of upstream wrapping and downstream contacts with the enzyme in open complexes under different regulatory conditions and promoter sequences. Also, we use transmission electron microscopy (TEM) and single particle analysis (SPA) to obtain a structural 3D model of a wrapped open complex assembled between RNAP and the wild-type λ*p*_R_ promoter. Finally, we used a real-time fluorescence assay to reveal the role of upstream and downstream DNA contacts in open complex formation and promoter escape.

## RESULTS

### Mechanical manipulation of single open complexes

Open complexes were formed at 37°C by mixing *E. coli* RNA polymerase-σ^70^ holoenzyme (RNAP-σ^70^) with a 2 kbp-long DNA construct bearing a single λ*p*_R_ promoter near its center. To select transcriptionally active open complexes, the mixture was incubated with heparin at 22°C, conditions known to destabilize other states in the initiation pathway of the open complex formation: RNAP + λ*p*_R_ ↔ RPc ↔ I_1E_ ↔ I_1L_ ↔ I_2_ → RPo, where RPc is the unwrapped closed complex and I_1E_, I_1L_, are wrapped closed complexes, and I_2_, is a wrapped open intermediate leading to formation of the stable open complex, RPo [1], [28]. AFM imaging confirmed that the enzyme binds near the center of the 2 kbp-long DNA where the λ*p*_R_ promoter is located (Figure S1, S2). A functional biochemical assay confirmed that open complexes were transcriptionally active (Figure S1A).

Next, we tethered open complexes between polystyrene beads and used optical tweezers to study them under force (Figure 1A). Force-extension curves obtained between 4 and 15 pN displayed a sudden, cooperative increase in length that we interpret as the mechanical disruption of promoter-RNAP interactions of an individual open complex. During the relaxation part of the experiment we also observed a cooperative shortening of the DNA template occurring at the same force than the extension. This observation indicates that the extension and contraction transitions occur near equilibrium. Surprisingly, this process could be repeated though more than 10 cycles at standard salt conditions (40 mM KCl) and in the rest of conditions tested (Figure 1B, additional examples are shown in Figure S12). Thus, promoter-RNAP contacts in open complexes could be mechanically disrupted and reformed reversibly. Overall, a total of N = 170 open complexes displaying n = 9484 transitions were analyzed. The percentage of molecules displaying transitions was about 50% for stabilizing conditions (e.g. glutamate) and about 10% for destabilizing conditions (e.g. high salt concentration); see Supplementary Table S1 for details. The remaining molecules that did not displayed transitions behaved mechanically as bare DNA. Recently, Cong *et al*., measured the strength of the enzyme with core promoter elements of the T1A1 promoter [29], however their experimental assay did not allow them to study the global role of upstream wrapping and downstream bending in transcription initiation.

**Figure 1.**
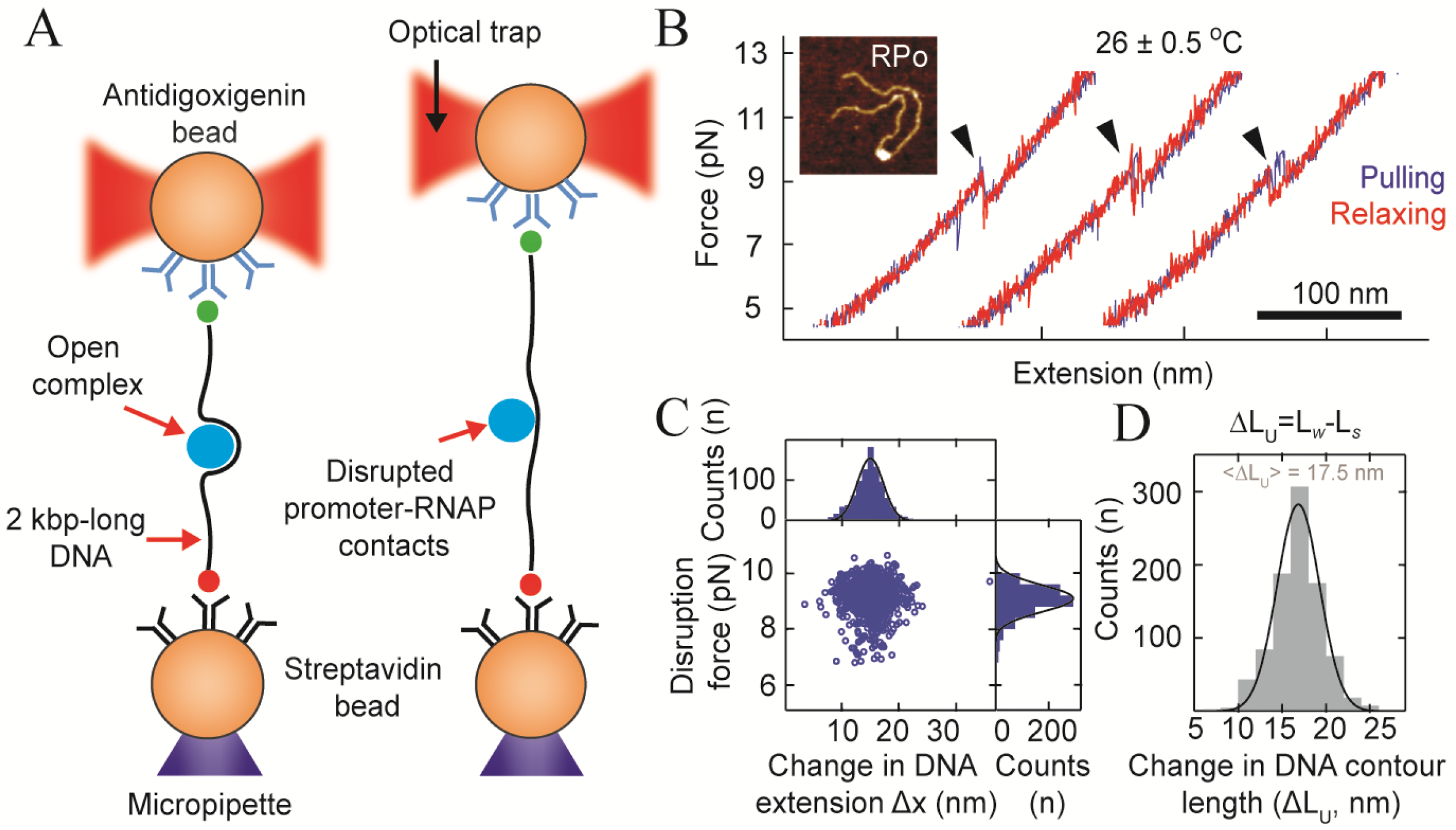
Single molecule experiment. **(A)** Optical tweezers experimental set-up: 2 kbp-long DNA bearing an *E. coli* RNA polymerase (RNAP, in blue) form an open complex near its center. One end of the DNA has biotin that is attached to the micropipette bead (orange) via a biotin/streptavidin bridge (red-black), while the other end has a digoxigenin, attached to an optical trap bead (orange) through a digoxigenin/anti-digoxigenin antibody association (green-blue). By moving the two beads relative to each other, pulling/relaxing cycles are applied to open complexes. **(B)** Representative force (in pN) vs extension (in nm) curves of three consecutive pulling (blue)/relaxing (red) cycles, between 4 and 12 pN at ~4 pN/s of loading rate. Scale bar: 100 nm; black arrows indicate the disruption (during pulling) and formation (during relaxing) or promoter-RNAP contacts, respectively. Inset: AFM micrograph of an open complex. **(C)** Disruption force (*F*_*U*_) vs change in extension (Δ*x*) plot and corresponding histograms distributions; continuous lines represent Gaussian fits. Statistics: ⟨*F*⟩ = 9.3 pN, σ_*F*_ = 0.6 pN, and ⟨Δ*x*⟩ = 15.5 nm, σ_Δ*x*_ = 3.2 nm. **(D)** Change in DNA contour length (Δ*L_U_*) histogram with ⟨Δ*L_U_*⟩ = 17.5 nm, 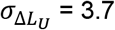 derived using the WLC model with a persistent length *P* = 20 nm. σ is the standard deviation of the mean.

75% of the open complexes assembled at the λ*p*_R_ wild-type promoter at standard salt conditions (40 mM KCl) displayed transitions at 8.8 ± 0.1 pN with no significant hysteresis and an average change in DNA extension, Δ*x*, of 15.5 ± 0.2 nm (Figure S1B-C, Tables S2, S3, S6, S7). Under similar salt conditions, about 25% of open complexes displayed two force-extension transitions. The first one occurred at 9.3 ± 0.2 pN, with a Δ*x* of 15.3 ± 0.4 nm and no hysteresis. The second one displayed significant hysteresis, with extension, Δ*x*, of 20.7 ± 0.1 nm occurring at 8.0 ± 0.1 pN, and a contraction, Δ*x*, of 18.4 ± 0.2 nm, occurring at 7.1 ± 0.1 pN (Figure S6, S7). The two apparent types of complexes—displaying either one or two transitions—were not interconvertible throughout the duration of multiple pulling/relaxing cycles (lasting from 10 to 90 cycles). Several control experiments revealed that the additional transitions observed in 25% of the molecules reflect the presence of non-specific DNA-protein complexes in the same tether (Figure S4-S7, Supplementary section 4). Therefore, we focused all further analysis on the reversible transitions occurring at ~9 pN.

### Promoter-RNAP interactions in open complexes are cooperative and reversible

Using the Worm-like Chain model of polymer elasticity [30], [31], the change in extension (Δ*x*) observed at ~9 pN corresponds to a DNA contour length increase *ΔL*_*rip*_ of ~17.5 nm, (Figures 1D, 3A, Table S4). This change in DNA contour length corresponds to the difference between the λ*p*_R_ promoter extension in interaction with the enzyme, *L_w_*, minus the distance along the pulling axis between the last two points of contact of the promoter around the protein (the secant in Figure 2A) given by,

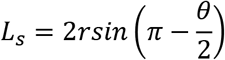

where the open complex has been approximated to a sphere of radius *r*, and where the promoter is assumed to bend by an angle *θ* around de enzyme. Then, *L_w_* = *θr* and, therefore, the change in DNA contour length upon the full straightening of the DNA during the disruption of promoter-RNAP contacts is,

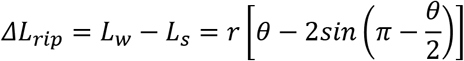

**Figure 2.**
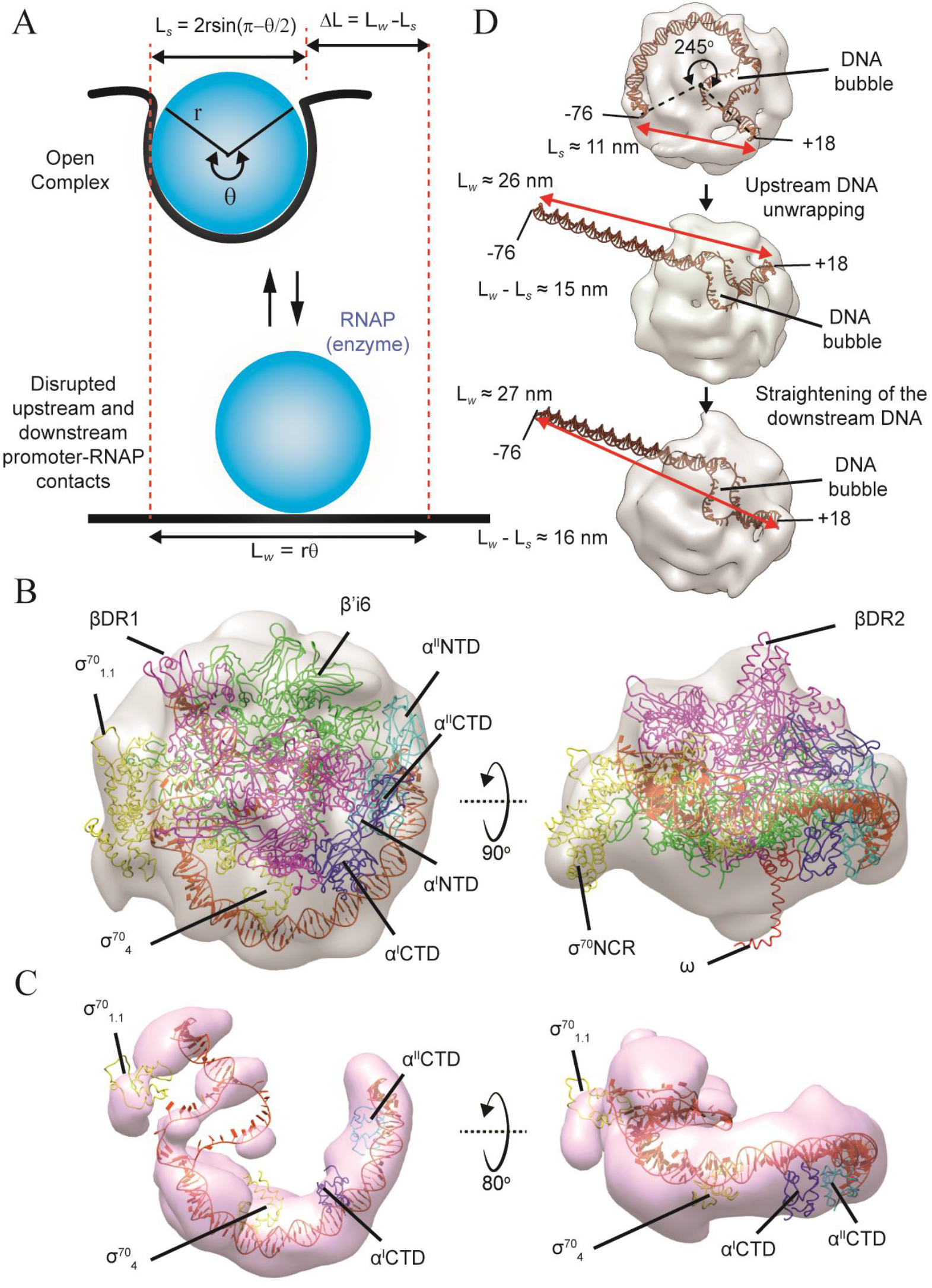
Open transcription initiation complex model. **(A)** Geometric model of the full λ*p*_R_ promoter interacting around the RNAP. **(B)** The 3D density map (gray surface, contoured at 0.42 σ) with the fitted pseudo-atomic model of the open complex. The β subunit is in magenta ribbon, βDR1 and βDR2 domains are pointed out. The σ^70^ subunit is in yellow ribbon, and their 1.1, 4 and NCR domains are indicated. The β’ subunit is in green ribbon and its β’i6 domain is pointed out. The αI and αII subunits are in blue and cyan ribbon respectively, with their N-terminal and C-terminal domains indicated. The omega subunit is shown as a red ribbon. The overall wrapped promoter is represented in orange ribbon with two blocks per base-pair. We use the “ hide-dust” tool of Chimera software to remove those densities not connected to the EM model at 0.42 σ. **(C)** Left: 3D difference-map is represented as a pink transparent surface contoured at 0.6 σ threshold level. This difference-map density depicts the path for wrapped promoter, extended from −76 to +18 positions (shown in orange ribbon with two blocks per base-pair), as well as the corresponding RNAP domains (respectively indicated) that were modeled to obtain the full open complex pseudo-atomic coordinates. Right: Rotated view of the difference-map. Densities not related to the DNA path were not shown for better visualization. **(D)** Top: distance between DNA ends (+18 and −76 positions) of the open complex is ~11 nm with an overall bent angle of ~245°. The angle was measured in projection view and the center of reference is the middle point of the longest dimension of the pseudo-atomic model (black point). Middle: representation of the partly unwrapped open complex, where the entire upstream DNA up to the transcription bubble is straightened. The distance between DNA ends (+18 and −76 positions) is ~26 nm. Bottom: representation of the fully unwrapped open complex, when the remainder DNA-downstream contacts are also disrupted. The distance between DNA ends is 27nm. The EM map is represented as a gray surface (density contoured at 0.7 σ) while the DNA is in orange ribbon representation.

Using *r* = 6 nm (assuming an RNAP radius of 5 nm plus half of the DNA width of 1 nm) and *ΔL*_*rip*_= 17.5 ± 0.3 nm, it yields *θ* = 4.5 ± 0.03 radians (257.9 ± 1.7°), *L_w_* = 27.0 ± 0.18 nm (~79 ± 0.52 bp), and *L_s_* = 9.5 ± 0.35 nm. The *L_w_* value is consistent with previous AFM studies [3], [7] and it was confirmed by our AFM measurements, which yielded a DNA compaction of 26 ± 2.5 nm (Figure S2A). Therefore, the single transition observed in pulling/relaxing experiments correspond to the disruption and straightening of the majority of promoter-RNAP contacts. Significantly, our results indicate that this process occurs in a two-state, cooperative manner, where no intermediates are observed.

Only the open complex and the I_2_ intermediate in λ*p*_R_ are long-lived complexes, due to the high energy barrier required to close the DNA bubble [32]. In contrast, the closed complex (RPc), I_1E_, and I_1L_ intermediates are short-lived because the DNA bubble is closed and a large energetic barrier does not protect them from dissociation. Therefore, the survival of promoter-RNAP contacts through several pulling/relaxing cycles (> 10), indicates that the complexes in our experiments are either open complexes or I_2_ intermediates. Moreover, it also indicates that the mechanical disruption of their interactions is not accompanied by the closing of the DNA bubble, which would lead to disassembly of the complex through its conversion to the short-lived intermediates RPc, I_1E_, or I_1L_. Roe *et al,* have shown that the open complex is significantly more stable than the I_2_ intermediate [33]. Thus, at equilibrium, the open complex population should represent > 99.9% of RNAP-λ*p*_R_ long-lived complexes. We remark that a previous single molecule study, using magnetic tweezers [34], detected and inferred the length of promoter-RNAP interactions in the open complex at the *lacCONS* promoter from the change in amplitude of transitions of negatively and positively supercoiled DNA. However, the limited spatial resolution of those experiments prevented further characterization of the mechanical disruption of promoter-RNAP contacts in the open complexes.

### Structural analysis of upstream promoter pathway in open complexes

To gain insight into their structure, open complexes were formed as described above but using a shorter DNA template (186 bp) harboring the λ*p*_R_ promoter. These complexes were deposited on a grid, negatively stained (NS), imaged by TEM, and subjected to SPA to obtain a 3D reconstruction at 17 Å resolution (Figures 2B, S8-S10, Table S8).

A difference-map, obtained by subtracting a 3D map generated from an X-ray structure of *E.coli* RNA polymerase holoenzyme alone (PDB 4YG2) [35] from our EM open complex model, shows a remaining density that depicts the path of the wild type λ*p*_R_ promoter in interaction with the enzyme (Figures 2C and S9C). This difference-map indicates that the λ*p*_R_ promoter bends around the enzyme with an overall angle of ~245° and extends from positions −76 to +18 along the template (Figure 2C, left), thus spanning a total length of 32 nm (Figure 2D, top). These results are in good agreement with: i) our estimated *L_w_* = 27 nm and *θ* = 4.5 radians (258°) values, and ii) footprinting studies that reveal an upstream periodic protection from the −35 to around the −70 position, and downstream protection up to around the +25 position [36]. Furthermore, this difference-map (Figure 2C) shows the densities corresponding to RNAP domains (α^I^CTD, α^II^CTD, σ^70^_1.1_ and σ^70^_4.1_) whose coordinates were missing in the starting template used for modeling (PDB 4YG2) (see materials and methods). We remark that previous structural studies reported upstream wrapping on transcription initiation complexes but in the presence of an activator protein and using promoters with an artificial transcription bubble [37, 38].

According to the EM model, the distance separating the last two points of contact of the λ*p*_R_ promoter with the enzyme is ~11 nm (Figure 2D, top), which is in good agreement with the *L_s_* = 9.2 nm value estimated from the optical tweezers experiments (Figure 2A). Then, if we assume that all upstream contacts down to the DNA bubble are mechanically disrupted by the force, the EM model would predict a contribution of *ΔL*_*rip*_ = *L_w_* − *L_s_* = 26 – 11 = 15 nm (Figure 2D, middle). Moreover, if DNA-protein interactions downstream of the DNA bubble are also disrupted, the straightening of the downstream DNA would contribute ~1 nm, making a total contour length change (*ΔL*_*rip*_) of 27-11 = 16 nm (Figure 2D, bottom). This value is close to the 17.5 nm obtained from our optical tweezers experiments (Figure 1D). Thus, the majority of the change in extension (Δ*x*) in the observed transitions (Figure 1B) corresponds to the promoter upstream wrapping, and a small fraction is likely due to promoter downstream bending in open complexes. Our results also indicate that these physically separated and independent DNA regions of the promoter are disrupted in a concerted and cooperative manner when subjected to force.

### RNAP alpha carboxy-terminal domains enable upstream wrapping

Upstream sequences of core λ*p*_R_ promoter consist of three upstream (UP) elements: proximal (positions −45 to −58), middle (positions −68 to −79), and distal (spanning from the −89 to −100 position) [27] (Figure 3B). AFM data suggest that UP elements interact with the carboxy-terminal domains of RNAP alpha subunits (α^I^CTD and α^II^CTD), and that they account for a significant fraction of upstream wrapping [21]. Interestingly, the 3D model shows that the promoter region around the proximal UP element experiences a sharp bending angle of ~115° (Figure 3C) that seems to interact with α^I^CTD (Figure 3D, left). This observation may explain the hypersensitivity to hydroxyl cleavage of λ*p*_R_ that has been described at around the −38 and −48 positions [36]. On the other hand, the region around the middle UP element adopts a smoother DNA bending angle of ~45° (Figure 3C) that appears to interact with α^II^CTD (Figure 3D, right). We do not observe a density corresponding to the distal UP-element in our EM model, suggesting that either this region does not contribute to the upstream wrapping of λ*p*_R_ in open complexes or that it only interacts transiently with the RNAP.

**Figure 3.**
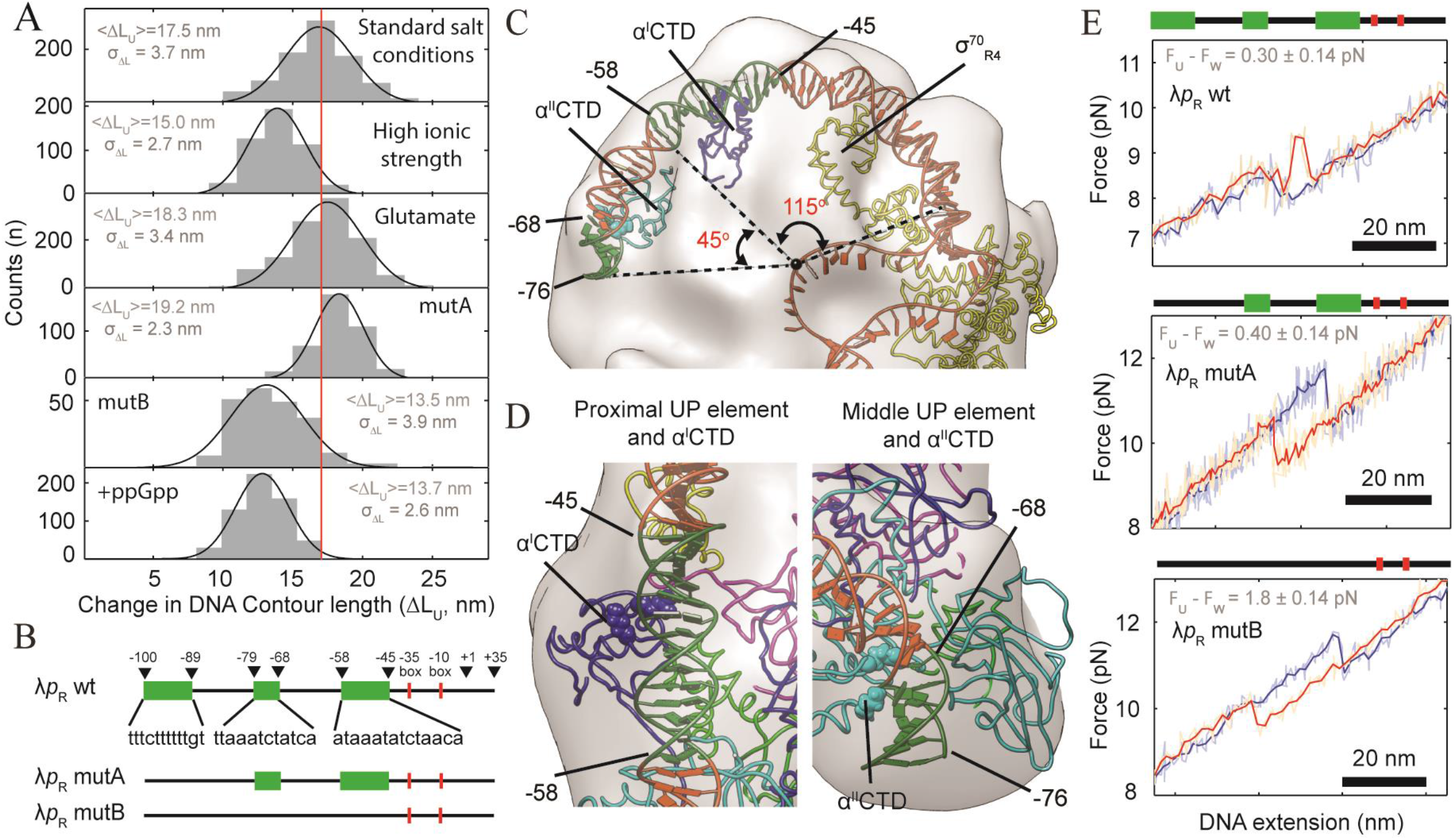
Role of upstream (UP) elements. **(A)** Δ*L*_*rip*_ histograms (gray) for all conditions tested. Black lines are the Gaussian fits to experimental data. The continuous red line across all graphs indicates the Δ*L_U_* mean value for λ*p*_R_ wild type at 40 mM KCl (standard salt conditions), ⟨Δ*L_U_*⟩ = 17.5 nm. (**B)** Diagram of the λ*p*_R_ wild type promoter from −100 to +35 positions. Upstream (UP) elements are in green, whereas −35 and −10 boxes are in red, mutA (lacking distal upstream, UP element), and mutB (lacking all UP elements). Black arrows and numbers mark positions on the promoter. **(C)** The open complex 3D density map (gray surface, contoured at 0.42 σ) and the fitted DNA promoter in orange color (ribbon) with UP elements in green. **(D)** Close-up view of the α^I^CTD, in blue ribbon, showing its possible interaction with the proximal UP element (left), and of the α^II^CTD, in cyan ribbon, with the middle UP element (right). These interactions would be mediated through some specific residues (spheres), as suggested by the crystal structure of CAP-αCTD-DNA complex [70] (see supplementary material). DNA and UP elements are represented as in (C). **(E)** Three examples depicting the trajectories of disruption (blue) and formation (red) of promoter-RNAP contacts of open complexes for the wild-type λ*p*_R_ (top), mutA (medium) and MutB (bottom) promoters. The difference between the transition force of disruption and formation of promoter-RNAO contacts (*F*_*rip*_ − *F*_*zip*_) provides a measure of hysteresis. Raw data (light color) was filtered and decimated to 100 Hz (darker color).

### Thermodynamics of promoter-RNAP contacts

Next, we determine the change in free energy, *Δ G*_*u* → *w*_, of promoter-RNAP interactions derived from the optical tweezers experiments. Because most force-extension transitions occur near equilibrium (Figure 1B), the free energy change associated with these interactions, *Δ G*_*u* → *w*_, is equal to the negative of the absolute value of the mean reversible work required to mechanically disrupt them, *W*_*w* → *u*_, (i.e., the area under the cooperative transition in the force-extension traces). After correcting for the entropic contribution of extending the DNA handles [38], we obtain the change in free energy at zero force 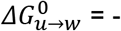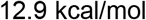 (Figure 4A-B, Table S5). Unexpectedly, this value is very close to the total change in free energy of the open complex formation (~−13 kcal/mol) determined in biochemical [32], [33] and AFM [3] studies under similar experimental conditions. Since we argue that the DNA bubble is not significantly affected by the application of force, this result indicates that the energy associated with the upstream wrapping and downstream rearrangements accounts for the majority of the open complex stability. Moreover, since the upstream DNA region in λ*p*_R_ stabilizes the open complex only by ~2 kcal/mol [22], we conclude that downstream rearrangements contribute the majority of the energetic stabilization of the open promoter complex, and that they are disrupted in a cooperative manner with the upstream wrapping interactions (Figure 4C).

**Figure 4.**
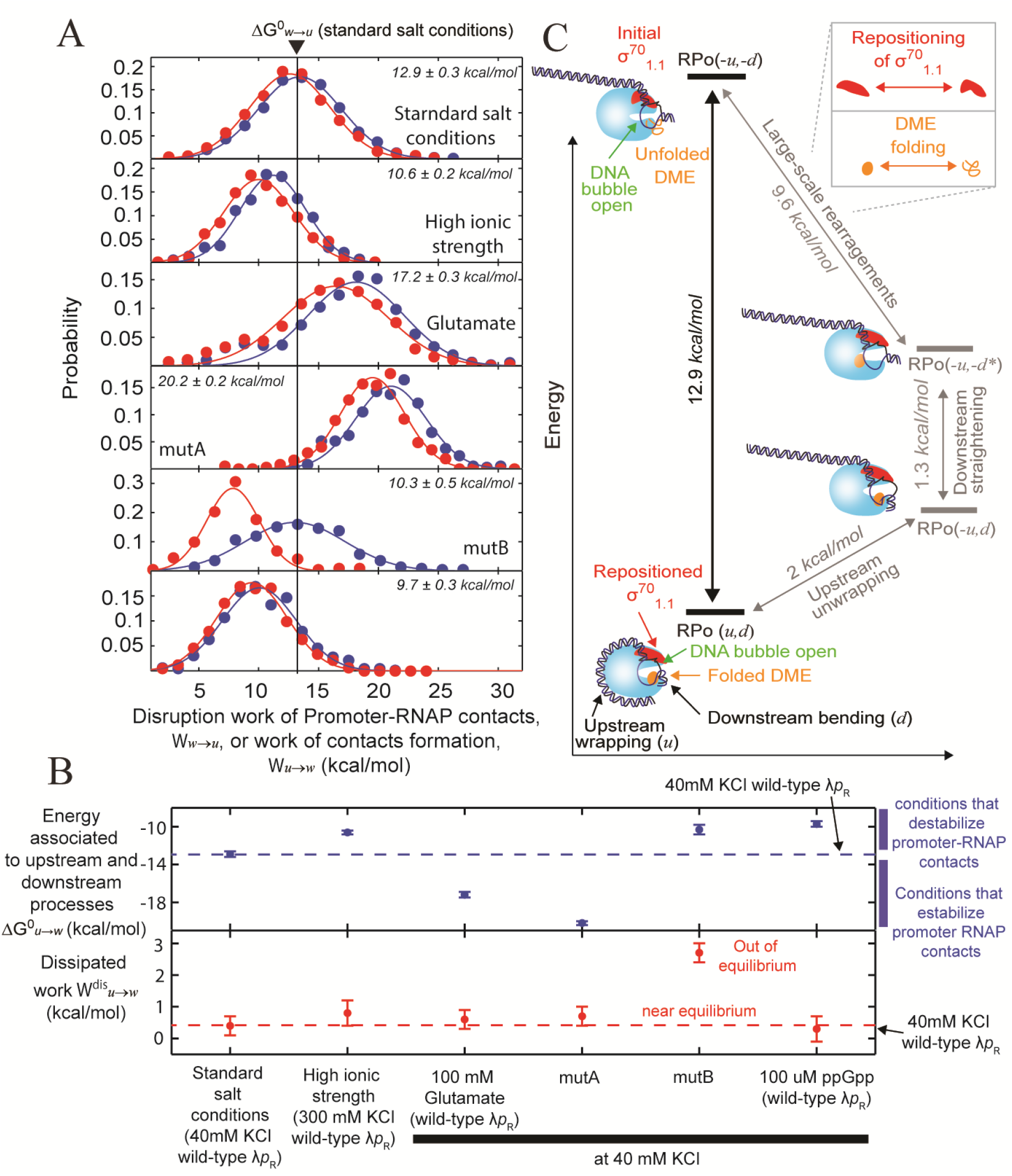
Energy associated with upstream wrapping and downstream rearrangement in open complexes. **(A)** Work distributions for disrupting (blue) and forming (red) upstream and downstream contacts. Lines and dots represent Gaussian fitting and experimental data of all conditions tested, respectively. Vertical black line across all graphs indicates the 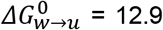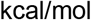 value for the λ*p*_R_ wild type at 40 mM KCl (standard salt conditions). **(B)** Top: 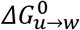 values for all conditions tested (kcal/mol). 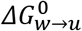 values larger than −12.9 kcal/mol indicate that the energy associated to upstream wrapping and downstream rearrangement is reduced (high ionic strength, in the presence of 100 μM ppGpp and the mutB construct). 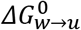 values smaller than −12.9 kcal/mol indicates that the energy associated with these processes has been increased (in the presence of glutamate and muA). Bottom: Dissipated work values for the mechanical disruption of these interactions 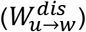. Only for the case of the MutB construct, the disruption and formation trajectories of upstream wrapping and downstream rearrangement are out of equilibrium. The dashed horizontal line represents λ*p*_R_ wild type at 40 mM KCl (top and bottom). **(C)** Thermodynamic cycle. The right vertical equilibrium (black continuous line) depicts the reversible and allosteric energetic coupling of upstream wrapping, downstream bending, and downstream large-scale rearrangements (DME folding and repositioning of the σ^70^_1.1_ domain) of open complexes under force in our optical tweezers experiments. Note that the DNA bubble (indicated in green) is not significantly affected. This process involves an overall free energy change 12.9 kcal/mol. The σ^70^ subunit is color coded red, σ^70^_1.1_ domain is the region that moves. Upstream wrapping and downstream bending are pointed out using black arrows. The downstream mobile elements in the β′-jaw (DME) is depicted in orange. Here, RPo (*u,d*) refers to a open complex that maintain upstream wrapping and all downstream interactions related to downstream bending and its large-scale rearrangements. RPo (*−u,d*) is the previous open complex but upstream interactions (wrapping) are missing; RPo (*−u,−d\**) is when the open complex is also missing the downstream bending but keeping the large-scale rearrangements. Finally, RPo (*−u,−d*) is the open complex missing all upstream and downstream contacts. The right bottom equilibrium (grey continuous line), refers to the upstream wrapping, involving ~2 kcal/mol [22]. The rigth middle equilibrium depicts the downstream DNA bending with an energy of ~1.3 kcal/mol. The rigth top equilibrium refers to the DME folding (orange) and the repositioning of the σ^70^_1.1_ domain (red) (inset), involving a residual energy of ~9.7 kcal/mol.

To validate our observations, we repeated pulling/relaxing protocols under open complex destabilizing (high salt concentration, 300 mM KCl) and stabilizing conditions (100 mM of potassium glutamate). As expected for electrostatic DNA-protein interactions, high salt concentration decreased the contribution of upstream wrapping and downstream rearrangements to open complex stability by ~+2.3 kcal/mol 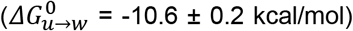, and reduced *Δ L*_*rip*_ by ~4 nm. In contrast, glutamate increases the energy associated with upstream wrapping and downstream rearrangements by ~−4.3 kcal/mol 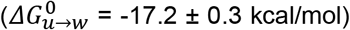, without variation in *ΔL*_*rip*_ (Figures 3A, 4A-B, Tables S4, S5). These results are in agreement with changes in open complex stability reported in bulk studies under similar conditions [39], [40]. Preliminary results with the unrelated Ampicillin (Bla) promoter indicate that the promoter-RNAP interactions in this case are also extensive, spontaneous, cooperative, reversible, and involve a large change in free energy (data not shown). These observations suggest that the energy associated with upstream wrapping and downstream rearrangements may be shared by other promoters.

### Role of upstream (UP) elements in promoter-RNAP interactions

To further characterize the contribution of upstream (UP) elements to the strength of promoter-RNAP contacts in open complexes, we first replaced the distal UP element by a random sequence, referred to as λ*p*_R_ mutA (Figure 3B). This sequence is identical to a construct previously used in an AFM study that showed a promoter wrapping length is reduced by ~9 nm [27]. Interestingly, we did not observe a reduction in *ΔL*_*rip*_ in our force-extension measurements (Figure 3A, Table S4). This result agrees with the EM model that does not show a density corresponding to the distal UP element in the open complex (Figure 3C). However, we obtain a 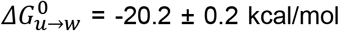, indicating that the distal UP element in the wild-type λ*p*_R_ promoter actually reduces the stabilizing energy of the upstream wrapping and downstream rearrangements by about +7.3 kcal/mol relative to the mutA construct (Figures 4A-B, Table S5).

Next, we replaced all UP elements by random sequences—construct λ*p*_R_ mutB (Figure 3B). Mangiarotti *et al*., reported that this mutant abolishes the majority of upstream promoter contacts in open complexes [27]. However, we detect a small fraction (~10%) of molecules displaying cooperative transitions and with a reduction of *ΔL*_*rip*_ of ~4 nm (Figure 3A). In these instances, we observe hysteresis between the disruption and re-formation of promoter-RNAP interactions, occurring at 10.5 and 8.7 pN, respectively (Figure 3E, Table S2). Thus, in the absence of the proximal and middle UP elements, upstream wrapping and downstream rearrangements occur out of equilibrium despite the fact that the pulling and relaxing rates were the same as those used with the wild type (Figure 3E, bottom). Therefore, for the mutB construct, the rate at which we perturb the system is faster than that at which upstream wrapping and downstream rearrangements can equilibrate. We conclude that the effect of UP elements is not only thermodynamic (stabilizing in the case of the proximal and middle elements and destabilizing in the case of the distal element), but also kinetic in the case of the proximal and middle elements, increasing the relaxation rate of the open complex in response to perturbations. Using the Crooks’ fluctuation theorem [41], we estimated a 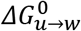 of −10.3 ± 0.5 kcal/mol for the mutB construct (Figure 4A-B). This observation suggests that interactions of RNAP with full upstream random DNA sequences can still enable wrapping and the energy associated to upstream wrapping and downstream rearrangements is ~80% of the energy of the wild-type λ*p*_R_ promoter. Overall, our results suggest that the stability conferred to the open complexes by the upstream wrapping and its coupled downstream rearrangements, does not depend on a specific upstream sequence (e.g. UP elements), but rather on the electrostatic and mechanical properties of the promoter in its interaction with the polymerase. Similarly, bulk studies showed that for *lacUV*5 and *gal*P1 promoters, upstream DNA-αCTDs interactions are not sequence specific [26], [42]. Recently, it was found that a long-lived open complex is also formed in the absence of UP elements in the *rrnB* P1 promoter although the number of such complexes is very small [43].

### ppGpp destabilizes promoter-RNAP interactions

The large energy associated with upstream wrapping and downstream rearrangements, suggests that modulation of these processes may play an important role in the regulation of DNA transcription. A recent AFM study has shown that the extent of upstream wrapping in open complexes of λ*p*_R_ is significantly reduced when RNAP-σ^70^ is pre-incubated with the transcriptional modulator ppGpp [3]. The reduced upstream wrapping correlates with a lower apparent affinity of the enzyme for the promoter in the presence of this metabolite [3]. The authors speculated that under environmental stress conditions, high levels of ppGpp decrease the extension of upstream wrapping and, therefore, the open complex stability at the λ*p*_R_ [3]. To further investigate the ppGpp mechanism, we repeated the force-extension experiments in the presence of saturating concentration of ppGpp (100 μM) to determine its effect on 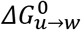 and *ΔL*_*rip*_. We still observe cooperative transitions of disruption/formation of promoter-RNAP contacts but with a decrease in *ΔL*_*rip*_ of ~4 nm and destabilization of the complexes of +3 kcal/mol 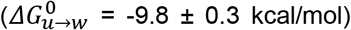 (Figures 4A-B, Table S4, S5). The decrease in the *ΔL*_*rip*_ value is smaller than that reported in AFM measurements (~8 nm) [3].

The fact that ppGpp destabilizes the complex (Figures 4A-B) more than the total amount of stabilization provided by the upstream wrapping alone (~2 kcal/mol, [22]), indicates that it also affects the downstream interactions and it lends support to the existence of an energetic coupling between the upstream wrapping and downstream rearrangements. The allosteric mechanism of ppGpp is suggested by two observations. First, our EM model shows that upstream promoter regions mainly interact with αCTDs, away from the ppGpp binding site, which is known to be located between the β’ and ω subunits [44]. Second, adding ppGpp at the concentration of 100 uM, corresponds to an increase in molar ionic strength of *ΔI* = 0.5×0.1×(4)^2^ = 0.8 mM, which is insignificant compared to the molar ionic strength of the standard salt conditions (40 mM KCl) used in our experiments, *I* = 0.5×40×(1)^2^ = 20 mM. This analysis suggests that ppGpp weakens the interactions between UP elements (proximal and middle) and the αCTDs, reducing the extent of upstream wrapping and allosterically causing a destabilization of upstream and downstream promoter-RNAP interactions.

### Upstream wrapping speeds up transcription bubble formation

Since upstream wrapping is allosterically coupled to downstream processes that represent the main stabilization of the complex, we wondered whether upstream wrapping plays a role in bubble formation and/or promoter escape. To this end, we used a bulk assay [45] to monitor the fluorescence of the λ*p*_R_ promoter (−100 to +18) containing a Cy3 label in position +2 (100-λ*p_R_*-Cy3 construct) (Figure 5A). In this assay, in the presence of 18-fold excess of RNAP, the Cy3 fluorescence increases initially fast reflecting the RNAP-λ*p*_R_ promoter association and then more slowly due to bubble formation. In these conditions, addition of a nucleotide mixture (NTPs) and heparin leads to promoter escape, which is accompanied by a decrease of Cy3 fluorescence [45] (Figure 5B). Analyses of the corresponding fluorescence vs time trajectories yield the effective rate constants of RNAP-λ*p*_R_ promoter association (*k_a_*), bubble formation (*k_o_*) and promoter escape (*k_e_*) [45] (Supplementary material, section 7, values are reported in Tables S10 and S11). We also compared the behavior of the full-length promoter (100-λ*p*_R_-Cy3) with that of a truncated λ*p*_R_ promoter (40-λ*p_R_*-Cy3) where all upstream sequences were removed (up to position −40), and no upstream wrapping is possible.

**Figure 5.**
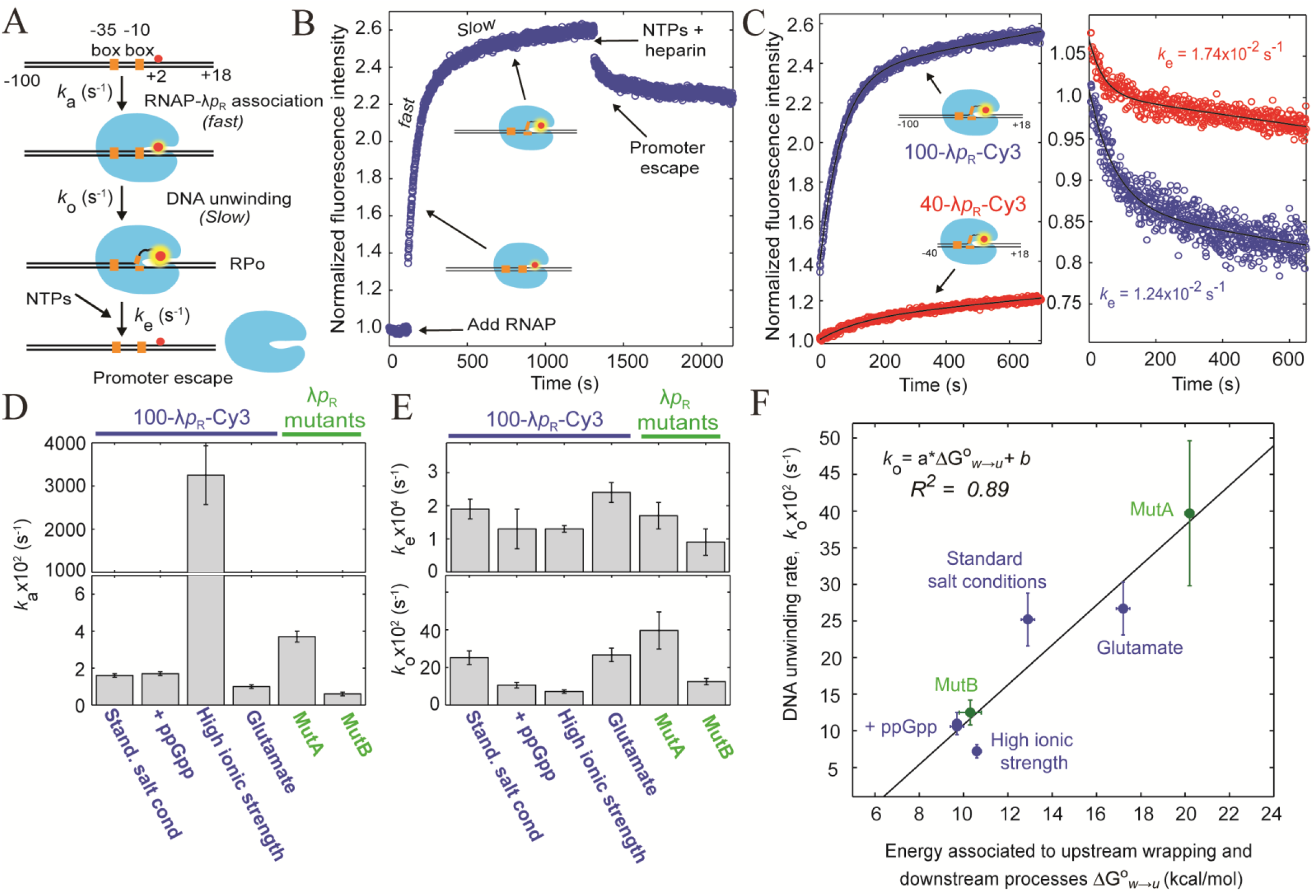
Real time open complex formation and promoter escape by fluorescence. **(A)** Experimental setup: Cy3 labeled (red dot) λ*p*_R_ promoter (DNA in black); −10 and −35 boxes (orange squares), and RNAP (light blue). Open complex formation/dissociation is described as a three-step process: RNAP-λ*p*_R_ association (*k_a_*), DNA bubble formation (*k*_*o*_) and promoter escape (*k*_*e*_) after addition of NTPs. **(B)** Fluorescence time course of a fully wrapped open complex (100-λ*p*R-Cy3). Cy3 fluorescence rapidly increases upon addition of 45 nM RNAP, it modestly increases during DNA opening and it decreases upon addition of 500 μM NTPs/heparin mix that induces the promoter escape. **(C)** Comparison of the kinetics of open complex formation/dissociation of a fully wrapped (blue trace) and unwrapped (40-λ*p*R-Cy3, red trace) complex; DNA bubble formation (left) and promoter escape (right), at low salt concentration (40 mM KCl). Black lines represent the two-exponential fitting. **(D) – (E)** The graphs compare the mean values of *k*_*a*_, *k*_*o*_, and *k*_*e*_ with their standard error of the Mean (SEM), obtained under stabilizing and destabilizing conditions tested **(F)** Positive linear correlation between the energy associated to upstream wrapping and downstream rearrangements 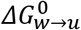 and the DNA bubble formation rate at different experimental conditions (left).

When wrapping is present (100-λ*p_R_*-Cy3), the kinetic analysis yields, *k_a_* ~1.6×10^−2^ s^−1^, *k*_*o*_~25×10^−4^ s^−1^ and *k*_*e*_ ~2×10^−2^ s^−1^ (Figure 5D-E). Truncation of the promoter’s UP elements (40-λ*p_R_*-Cy3) visibly affects the fluorescence profile (Figure 5C), with a six-fold decreased of *k*_*o*_ (*p* = 0.009) and not significant changes in *k*_*e*_ and *k*_*a*_ (Figures 5D-E and Table S11). These results indicate that upstream wrapping facilitates bubble formation, in agreement with previous observations using nitrocellulose filter-binding assays, in which faster open complex formation is observed in the presence of the upstream regions for λ*p_R_* and *lacUV*5 promoters [22], [26].

### The energy associated with promoter-RNAP interactions correlates positively with the rate of bubble formation

Next, we performed fluorescence experiments of the 100-λ*p_R_*-Cy3 construct under open complexes stabilizing (100 mM glutamate) and destabilizing (300 mM KCl, 100 μM ppGpp) conditions. Corresponding values of *k*_*a*_, *k*_*o*_ and *k*_*e*_ are shown in Figures 5D-E and Table S11. First, conditions that change the energy of upstream wrapping and downstream rearrangements, alter RNAP-λ*p_R_* association rate, but generally they are not correlated (Table S11). Second, we find that the strength of these promoter-RNAP contacts with the enzyme correlates positively with the bubble formation rates (Figures 5E-F, left). Formation of the transcription bubble (I_1L_ ↔ I_2_) is the rate limiting step of the open complex formation [1], [32]. Therefore, this result suggests that the energy associated with upstream wrapping and downstream rearrangements affect the rate of open complex formation and that early steps of the complex formation should be sensitive to tension. Third, it is known that mutations in the core sequences (at the −10 and −35 boxes), longer discriminators (the region between −10 and +1), and mutations in the downstream DNA region, facilitate promoter escape [45]–[48]. While it is usually assumed that the stronger the promoter-RNAP contacts, the slower the promoter escape should be, those studies do not determine the effect of such modifications on the energy of promoter-RNAP contacts. Here, we do not find a strong correlation between the energy associated to upstream wrapping and downstream rearrangements and promoter escape (Figure 5E-F, right).

To determine if the correlation between the strength of promoter-RNAP contacts with the rate of bubble formation (*k*_*o*_) is affected by the presence or the absence of UP elements in the promoter, we performed experiments using λ*p*_R_ mutant versions that destabilize (100-MutB-Cy3) or stabilize (100-mutA-Cy3) open complexes at standard salt conditions (Figure 4B). Interestingly, the correlation observed above between the energy associated with upstream wrapping and downstream rearrangements and the rate of bubble formation is retained with these λ*p*_R_ mutant variants (Figure 5F). Similarly, only a weak correlation is observed between the values of promoter scape rates and the changes in stability associated with these mutants (Figure 5F). Altogether these results and those obtained in the presence of glutamate, high salt concentration, and ppGpp, strongly suggest a regulatory role of upstream wrapping on downstream rearrangements and on the rate of bubble formation.

### Upstream wrapping reduces the effect of ppGpp on bubble formation rate

It has been proposed that ppGpp has a negative allosteric effect on bubble formation by modulating the β’-clamp dynamics [49]. As illustrated in Figures 5D-F, our results indeed confirm that ppGpp slows down *k*_*o*_ by ~50% (*p* = 0.02) without affecting RNAP-λ*p*_R_ promoter association (*p* = 0.52) or promoter escape (*p* = 0.41) (Figure 5E and Table S11), a result consistent with previous bulk studies [50], [51]. Therefore, ppGpp might have an allosteric dual role on transcription initiation: i) it reduces the thermodynamic contribution of upstream wrapping and downstream rearrangements to open complex stability; and ii) it slows down bubble formation. Interestingly, in the absence of upstream wrapping (40-λ*p_R_*-Cy3), ppGpp slows down even more (~eight-fold) the rate of bubble formation (Table S10). Thus, upstream wrapping reduces the effect of ppGpp on the rate of bubble formation. This inference helps to explain *in-vitro* and *in-vivo* observations [51]–[53] that promoters forming short-lived open complexes (presumably due to weak promoter-RNAP contacts), such *rrnD* P1 and *rrnB* P1 [3], are strongly inhibited by ppGpp [51]; whereas promoters forming long-lived open complexes, such as λ*p*_R_, characterized by extended upstream wrapping [7], and *lacUV5* promoters [13], are more resistant to ppGpp.

## DISCUSSION

### Upstream wrapping, downstream bending and its associated large-scale rearrangements are energetically coupled to form the open complex

The open complex formation involves a number of steps: RNAP + λ*p*_R_ ↔ RPc ↔ I_1E_ ↔ I_1L_ ↔ I_2_ → RPo. Bulk studies suggest that the last step of open complex formation (I_2_ ↔ RPo) makes the largest energetic contribution to the stability of the complex [1], [6], [28] (Figure 6A). This step involves the accommodation of downstream DNA within the cleft formed by the β′-jaw/β’-clamp domains of RNAP, the folding of downstream mobile elements in the β′-jaw (DME), and the repositioning of the σ^70^_1.1_ domain [1], [6], [28]. In our single molecule experiments upstream wrapping and downstream bending display a change in DNA contour length of about 17 nm and involve a total energy of around −13 kcal/mol (Figure 4A-B). The mechanical disruption of this process in open complexes occurs at around 9.3 pN. The straightening of the downstream DNA bend is expected to contribute only ~1 nm to the change in extension (*Δ x*) observed in the optical tweezers experiments (Figure 2B). Thus, we can estimate that its energetic contribution would be of about −9.3 pN.nm or ~−1.3 kcal/mol to the open complex stability (Figure 4C). That leaves a residual stabilization energy of −11.7 kcal/mol. On the other hand, Davis *et al*. have shown that the energy associated with wrapping of the upstream promoter segment contributes only ~−2 kcal/mol in λ*p_R_* [22]. Accordingly, the upstream wrapping and downstream DNA bending contribute a total energy of ~ 3.3 kcal/mol (Figure 4C). This result is consistent with a recent study using bulk FRET experiments [23]. The difference of ~−9.7 kcal/mol must correspond to large-scale rearrangement associated to downstream bending which represent the largest contribution to the open complex stabilization (Figure 4C).

**Figure 6:**
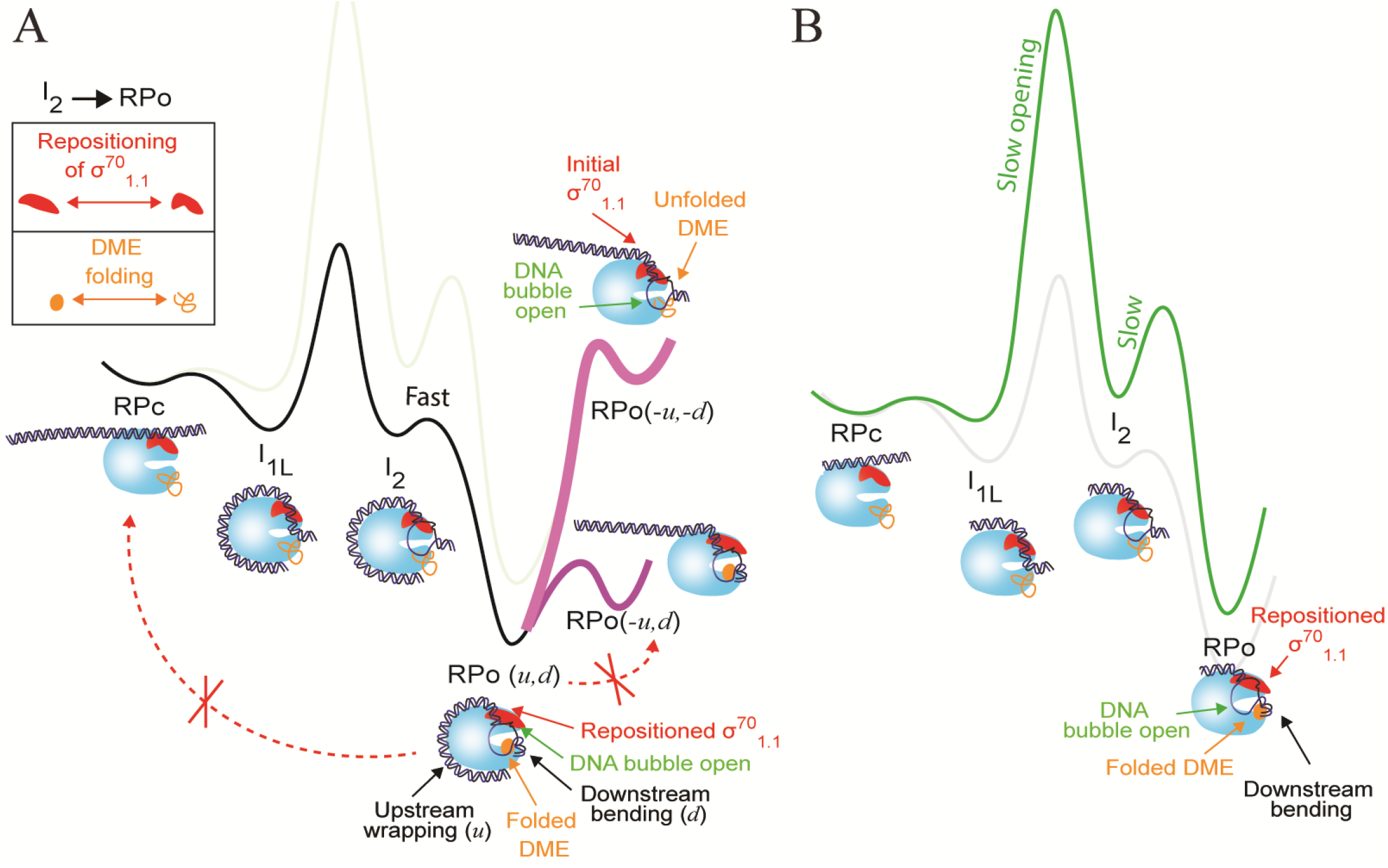
Role of upstream wrapping and downstream rearrangements in open complex formation. **(A)** Thermodynamic model of open complex formation for the full λ*p*_R_ promoter (when wrapping exist). The closed complex (RPc), the fully wrapped (I_1L_), the open intermediate (I_2_), and the final open complex (RPo) are indicated. The change in free energy due to upstream wrapping and downstream bending (RPc → I_1L_) is ~−2 kcal/mol [22], [23] and ~1.3 kcal/mol (see text), respectively; DNA bubble opening (I_1L_ ↔ I_2_) has a large but unknown activation energy [32], large-scale rearrangements (DME folding and repositioning of the σ^70^_1.1_ domain) stabilize the open complex by ~−9.6 kcal/mol during the late step (I_2_ → RPo) [33], and the overall change in free energy of the open complex formation is ~−13 kcal/mol [3], [32], [33]. The mechanical transition induced in the present study is indicated in pink between the final open complex, RPo (*u,d*) and an open complex missing upstream and downstream contacts, RPo (*−u,−d*). The dotted red arrow with the x mark (left) indicates that the mechanical disruption of upstream (*−u*) and downstream (*−d*) interactions to form the close complex (RPc) is not allow in our experiments since the closing of the DNA bubble involves an enormous energy barrier. Also, the dotted red arrow with the x mark (rigth) indicates that the mechanical manipulation of open complex cannot disrupt only upstream contacts to form RPo (*−u,d*). **(B)** Thermodynamic model of open complex formation for a truncated λpR promoter up to position −40 (no upstream wrapping is possible). The Energy landscape **(A)** and **(B)** were modified from Ruff *et al*., [1].

Surprisingly, the mechanical disruption experiments presented here, indicate that in the open complexes, upstream wrapping, downstream bending and its large-scale rearrangements occur as a single, cooperative, and reversible transition, indicating a high degree of energetic coupling between these processes (Figure 4C). If this were not coupled, we should have observed two transitions occurring in series as we pull the complex in the optical tweezers. First, we would have seen a large transition of 16 nm corresponding only to upstream unwrapping, occurring at a force of approximately 0.9 pN since 0.9 pNx16 nm = 14 pN.nm ≈ 2 kcal/mol. As we continue pulling, we would have observed a small 1 nm transition occurring at a force above 70 pN, since the residual energy of downstream bending with its associated large-scale rearrangements is 11 kcal/mol ~76 pN.nm. The existence of upstream and downstream allosteric coupling in initiation complexes is supported by: i) a structural study showing that upstream rearrangements in elongation complexes, induced by an RNA hairpin, generate conformational changes that propagate to the downstream region [54]; and ii) the finding that mutations in the upstream region in λ*p*_R_ and *lacUV*5 promoters, modify the protection of the downstream promoter by the RNAP [22], [26], [55]. Thus, our observed transitions induced by the application of force in the open complex, correspond to the cooperative and simultaneous unwrapping of the upstream DNA, the straightening of the downstream DNA, and their associated large-scale downstream rearrangements.

Note that upstream wrapping and downstream bending also occur in the close complex [23], whereas the large-scale rearrangements associated to downstream bending have been proposed to occur after the opening of the DNA bubble [1], [6], [28]. Our results indicate that the upstream wrapping and the downstream rearrangements are energetic coupled and occur in a concerted manner with DNA bubble formation during the formation of the open complex. Note that the allosteric coupling between these spatially separated processes is both thermodynamic (stabilizing) and kinetic in nature (Figure 6A). Indeed, removal of the upstream UP elements (which eliminates wrapping) does not prevent the ensuing of the downstream promoter processes and their stabilizing effects on the complex [22, 23], but most likely affects the rate at which these downstream processes concomitant with DNA bubble formation occur (Figure 6B). This inference could provide an explanation to our observation that the energy associated with the overall mechanical disruption of upstream and downstream contact with the enzyme correlates positively with the rate of DNA bubble opening (Figure 5F). Interestingly, the repeatability of the pulling/relaxation cycles in our mechanical experiments, indicate that the disruption of these upstream and downstream contacts does not lead to the closing of the DNA bubble (Figure 6A), which is likely the result of the large energetic barrier known to be associated with this process [32]. It is interesting to speculate that the upstream and downstream allosteric coupling established in the open promoter complex may be preserved in actively transcribing elongation complexes [54].

### Biological Consequences

Upstream wrapping and downstream rearrangements are conserved phenomena in open complexes across species, from prokaryotic to mammalian cells [7]–[15][16]–[20]. Here we have shown that these processes besides providing the largest contributions to the stabilization of the open promoter complex, are allosterically and reversibly coupled, suggesting the existence of an intra-molecular signaling pathway between these two regions. We find that the strength of these promoter-RNAP contacts in the open complex are positively correlated with the rate of DNA bubble formation. Moreover, we have shown that upstream and downstream events are enabled by αCTD interactions with upstream DNA, irrespective of its sequence, suggesting that its thermodynamic consequences in the open complex stability are intrinsic to the enzyme and the polyelectrolyte properties of the DNA promoter. Finally, our results suggest that the coupling between upstream and downstream events are part of a c*is*-regulatory network, that could furnish a mechanism of control of gene expression by protein factors and regulatory metabolites. Any modulator that alters the thermodynamic contribution of this coupling, or affect the velocity of the DNA bubble formation, such as upstream-binding transcriptional factors, CAP, HN-S, Hu, FIS, or downstream-binding factors such Gp2, could finely tune gene expression in response to the needs of the cell.

## Materials and Methods

### Transcription activity of open complexes

We assembled complexes by mixing 500 nM of RNAP (Kashlev laboratory, National Institutes of Health, Bethesda, MD, USA) and 100 nM of 231 bp-long DNA construct, bearing a λ*p*_R_ promoter sequence in the center, in TB40 buffer (20 mM Tris-HCl pH 7.9, 40 mM KCl, 5 mM MgCl_2_, 1 mM DTT), and incubated the mixture at 37°C for 20 min [7], [36]. To favor the selection of specific open complexes, we added heparin to the mixture to a final concentration of 200 ng/ul and incubated it at room temperature for 15 min [40]. The transcription reaction was started by adding [α-^32^P]UTP (1 μCi; 1 Ci = 37 GBq). After 10 min, we added NTPs to a final concentration of 0.5 mM ATP, CTP, GTP, UTP (Sigma-Aldrich, USA) and incubated the mixture at room temperature. Later, we collected aliquots after 0, 10, 20, and 30 sec of incubation and stopped each reaction by mixing it with loading buffer supplemented with formamide. We visualized the transcription products by electrophoresis in a denaturant gel (23% polyacrylamide, 7 M urea) run at 800V for 2 h (Bio-rad, Hercules, CA, USA).

### AFM imaging of open complexes

We assembled open complexes as described above but using 20 nM of a 2 kbp-long DNA, 60 nM RNAP and 2 ng/ul of heparin. The reaction was diluted down 20 times in TB40 and 2 ul were deposited onto freshly cleaved mica for 2 min, rinsed with water and dried with nitrogen flow. AFM imaging was carried out in air using tapping mode in a multimode Nanoscope 8 microscope (Bruker, Billerica, MA, USA). Images of 512×512 pixels were collected with a scan size of 2 × 2 μm at a scan rate of 1.5 lines per second. We measured the DNA contour length of intact free DNA molecules and open complex as described in [3]. Algorithms are described elswhere [56], [57]. Complexes with RNAP bound to the DNA ends and with more than one RNAP were discarded from this analysis.

### Preparation of open complexes for optical tweezers studies

We produced a 2 kbp-long DNA bearing the λ*p*_R_ promoter at the center by PCR, using a digoxigenin-labeled forward primer and a biotin-labeled reverse primer (Integrated DNA Technologies, Coralville, IA, USA), and Phusion DNA polymerase (New England Biolabs, Ipswich, MA USA). DNA product was gel purified. We assembled open complexes as described above but using 1 pmol of RNAP and 0.2 pmol of DNA in TB40. The reaction was then diluted 20 times in TB40. Later, 3 μl of diluted open complex-heparin solution were incubated with 5 μl of 3.4 μm anti-dig beads for 15 min at room temperature. The immobilized open complex solution was diluted in 500 μl of TB40 before pulling/relaxing experiments in the optical tweezers. We performed most of our experiments in 40 mM KCl (TB40), we also refer to this condition as the low salt concentration or standard condition. For high salt concentration, we used 300 mM KCl and in some experiments the complexes were incubated with 100 mM potassium glutamate (KGlu). Because the open complex lifetime is ~6 h [58], the samples were used only up to 3 h.

### Optical tweezers data collection

Single molecule manipulation of open complexes was carried out using a counter propagating optical tweezers instrument with 850 nm laser. We performed pulling/relaxing protocols of single open complexes [59]–[61] at 26 ± 0.5°C, at a loading rate of about 4 pN/s between 4 and 15 pN of tension, with a waiting time of 2 sec at the end of each cycle. The data was collected at 1 kHz. Several pulling/relaxation cycles were recorded per molecule during each experiment; lasting between 5 – 20 min. Single open complex tethers were identified when their rupture occurred in a single step and with the force dropping to 0 pN. Only pulling/relaxing traces displaying single tethers were selected for analysis.

### Optical tweezers data analysis

We estimated the change in the DNA contour length (*ΔL*_*rip*_) of the transitions using the Worm-like Chain (WLC) model of polymer elasticity [30], [31]. We calculated the reversible and non-reversible work as the area under each transition according to 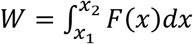 where *x*_2_ − *x*_1_ = Δ*x* is the change in DNA extension (nm) upon disrupting promoter-RNAP interactions at the force *F*(*x*_1_) [38], [41], [59]. For non-reversible transitions, we estimated the reversible work or free energy change (*Δ G*_*u*→*w*_) by using the non-reversible work distributions for disrupting (*W*_*w*→*u*_) and forming (*W*_*u*→*w*_) these promoter-RNAP interactions and the Crooks fluctuation theorem [41]. To obtain the free energy change at zero force 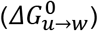, we subtracted the contribution arising from the entropy loss due to the stretching of the λ*p*_R_ promoter and of the DNA handles on both sides to the promoter (*W*_*strech*_ in Table S5) from *Δ G*_*u*→*w*_ [38], [41]. Pulling/relaxing trajectories were analyzed using a Matlab program developed in Bustamante’s laboratory. Work, transition force, change in DNA extension, and contour length distributions were fitted to a Gaussian distribution. Mean, standard deviation and error values (confidence interval at α = 0.05 of significance level of two sides), are reported per condition.

### Assembly of open complexes for TEM imaging

The open complex was assembled and selected as described above but using 60 nM 186 bp-long DNA and 30 nM of RNAP. This 1:2 molar ratio of RNAP-DNA was used in order to minimize the presence of free protein. No further purification was required. Then, we deposited 3 μL of the sample on an ultra-thin carbon-coated grid, negatively stained (NS) with 2% (m/v) uranyl acetate, and air-dried it at room temperature. We performed the data acquisition of 500 images in a JEM-2100 microscope (Jeol, Tokyo, Japan), equipped with an F-416 CMOS camera (TVIPS, Gauting, Germany) at 200 kV, with a pixel size of 1.78 Å and with a defocus range between −1 and −2 μm. In EM micrographs, free DNA and open complexes are clearly distinguishable (Figure S8).

### Single Particle Analysis (SPA)

We performed SPA of the raw data using IMAGIC-4D [62] software (Image Science, Berlin, Germany), following the most recent methodology [63], [64]. The final three-dimensional reconstruction was obtained from 16,015 particles classified into 666 class-averages. Figure S11A shows the open complex reconstruction contoured at 0.42 σ level. We were able to assign the densities corresponding to: i) the β dispensable region 1 (βDR1, residues 225–343, also known as β insert 4), ii) the β dispensable region 2 (βDR2, residues 938-1042, also known as β insert 9), iii) the β’ insert 6 (β’i6, residues 942–1129, also known as β’ trigger-loop non-conserved domains or β’GNCD), iv) the σ^70^ non-conserved region (σ^70^NCR), and v) the ω subunit [5], [27], [35], [65] (Figure S9A). The Fourier Shell Correlation (FSC) [66] of the EM reconstruction shows an estimated resolution of 17 Å, with 1/2-bit threshold criterion [67] (Figure S9B). The final open complex EM reconstruction was coarsened with 2-fold pixel binning and low-pass filtered according to the resolution estimated by the FSC. Density map have been deposited at EMDB with accession code EMD-0340, and it is also provided in Supplementary data set (3d_final_map.mrc).

### Generation of the open complex coordinate model

The pseudo-atomic coordinates of the open complex were obtained preserving the overall features of the crystallographic coordinates from PDB 4YLN [5] (structure that lacks the αCTDs components and DNA wrapping) and, through the process of modeling and optimization that we describe here for the wrapped DNA and its interface with RNAP. First, the crystallographic coordinates of the *E. coli* RNAP and transcription bubble DNA from [5] were manually fitted into the EM reconstruction using the UCSF Chimera software [68]. Second, the coordinates of the DNA region upstream the transcription bubble were modeled with the on-line server 3D-DART [69] and fitted in the remaining density of the EM reconstruction. Third, the modeling of the promoter region downstream the transcription bubble had to be performed with a different strategy. Since the PDB template contains the transcription bubble coordinates up to the downstream +12 position, we modeled a small promoter fragment from that position to the boundary of the density map. All this process was done following the unique possible and rationale direction, which is projecting this promoter region through the RNAP main channel. Using this modeling process, we reach the +18 position (6 bp more than the downstream end of the template), which we consider the downstream end for the wrapped DNA. Fourth, the αCTDs coordinates were obtained according to crystallographic structure PDB 1LB2 of the ternary complex CAP-DNA-αCTD [70]. The DNA in contact with αCTD present in PDB 1LB2 was semi-automatically aligned with our modeled DNA by translating and rotating it. The interface between the αCTD and our modeled DNA was optimized through energy minimization. This process, which was performed for each of the αCTDs, positioned these domains relative to the rest of the protein. At the end of this process, each αCTD appeared ~45 Å away from its respective NTD, a distance consistent with the 16-amino acid α-linker (residues 233-249) connecting both domains [35]. Then, the α-linker for each α subunit was modeled and optimized by energy minimization. Finally, the coordinates for the σ^70^_1.1_ domain (residues 9-89) were obtained [71], [72] and fitted into the EM density map. Also, the missing residues of this domain as well as those that connect it with the remaining portion of σ^70^ subunit were modeled and optimized according to their amino acid sequence [73]. All processes of DNA optimization and the energy minimization of the final full coordinate model were performed using the AMBER 99 force field available in the YASARA software [74]. Refined pseudo-atomic model has been deposited in the Protein Data Bank with accession code 6N4C, and it is also provided in Supplementary data set (RPfinal20.pdb).

To get a clearer visualization of the complete promoter path, a difference-map was generated between our EM map and a 3D density obtained from a previous published x-ray structure of *E. coli* RNAP holoenzyme [35]. The maps were aligned, filtered at the same spatial frequency and normalized before calculating the difference-map. All these processes were performed using IMAGIC-4D software [62]. This difference-map shows densities that can be assigned to the transcription bubble, upstream and downstream-DNA regions, as well as those RNAP domains not present in the initial template (PDB 4YLN).

### Real time fluorescence assay

DNA promoters for fluorescence transcription experiments were constructed by annealing and ligation of oligonucleotides (Integrated DNA Technologies, Coralville, IA, USA) with single-strand overhangs as described [45]. Each construct is synthesized using three double stranded oligos. For the mutA-Cy3construct we use the following oligonucleotides: mutA-sense/mutA-antisense, startUP-sense/startUP-antisense and downstream-sense^Cy3^/downstream-antisense. For mutB-Cy3: mutB^II^-sense/mutB^II^-antisense, mutB^I^-sense/mutB^I^-antisense and downstream-sense^Cy3^/downstream-antisense. For the 100-λ*p_R_*-Cy3: endUP-sense/endUP-antisense, startUP-sense/startUP-antisense and downstream-sense^Cy3^/downstream-antisense. For 40-λ*p_R_*-Cy3: upstream-sense/upstream-antisense and downstream-sense^Cy3^/downstream-antisense (Table S9). The downstream-sense (amino modifier C6 dT) was labeled with Cy3-NHS ester (Lumiprobe, Hunt Valley, MD, USA) at +2 position of the non-template strand (downstream-senseCy3). Labeling reaction was done in 0.2 M Na2CO3 pH 8.5, 10% DMSO, with shaking at 4°C for 24 h, and using a ~30-fold molar excess of Cy3 compared to the oligonucleotide. Free dye was removed via dialysis and the degree of labeling was ~90%. Annealed oligonucleotides were purified by PAGE and electroelution. Oligo duplexes were ligated in equimolar concentrations using T4 enzyme (Bio-rad, Hercules, CA, USA) for 2 h. Final DNA template was purified by PAGE and electroelution.

For fluorescence measurements, we recorded Cy3 emission at 570 nm, whereas excitation was at 550 nm on a Felix fluorometer (Horiba, Kyoto, Japan) at room temperature. First, we monitor fluorescence of 2.5 nM DNA construct containing a Cy3 molecule for ~3 min. Second, we added manually (mixing time was ~ 4 sec) *E. coli* RNAP (Bio-rad, Hercules, CA, USA) to a final concentration of 45.5 nM. The final promoter/RNAP ratio was ~1:18.2. We monitored the increase of Cy3 fluorescence (RNAP-λ*p*_R_ association and DNA bubble formation) until a plateou, which indicates open complexes were formed [45]. Finally, we manually added a NTP mixture (rCTP, rUTP, rGTP and rATP) and heparin to a final concentration of 500 μM and 34 mg/ml, respectively. We continue monitoring the Cy3 decrease of fluorescence (promoter escape) for ~15 min. For the case of 100-λ*p*_R_-Cy3 at high salt conditions, mutB-Cy3 at standard salt conditions and experiments with 40-λ*p*_R_-Cy3 we followed the same protocol but we monitored Cy3 increase of fluorescence for ~1 h, and we added heparin to a final concentration of 0.68 mg/ml. The method to extract the RNAP-λ*p*_R_ association (*k_a_*), the DNA bubble formation (*k*_*o*_), and the promoter escape rates (*k*_*e*_) are described elsewhere [45]. Rate constants are reported as mean value ± standard error of the mean (SEM) and fitting was performed using a confidence level of 95% (α = 0.05); and the statistical significance between two rate constants was evaluated using the two-tailed Student’s test (p < 0.05). Fluorescence data was analyzed with a non-linear fitting program code wrote in python 3.5. The source python codes are provided as supplementary data (binding-open.py and escape.py).

## Supporting information

SUPPLEMENTARY MATERIALS, METHODS and DISCUSSION

## Acknowledgments

We thank Prof. Marin van Heel, Prof. Claudio Rivetti and Bustamante laboratory members for fruitful discussions. We thank Steven Smith for assistance with the optical tweezers device (minitweezers), Dr. Mikhail Kashlev for providing the RNAP polymerase enzyme for preliminary experiments, and John Van Patten, Cristhian Cañari and Piere Rodríguez for critical reading of the manuscript. We also thank Omar Herrera for assistance in the SDS transcription activity, Alexander Robles for his assistance to write the program in Python 3.5 to analyze fluorescence data and LNNano/CNPEM for the access to the EM facility. This research was supported by the MSc Scholarship from Consejo Nacional de Ciencia Tecnologia e Innovacion tecnológica (CONCYTEC) [code 014-2013-FONDECYT] to [R.P.S.]; and MSc scholarship from the Coordenação de Aperfeiçoamento de Pessoal de Nível Superior-Brasil (CAPES) [code 001/88882.143467/2016-01] to [A.J.F.A.]. This work was also supported by National Institutes of Health (NIH) [grant number 01GM032543] to [C.B]; the US Department of Energy Office of Basic Energy Sciences Nanomachine Program under [contract DE-AC02-05CH11231] to [C.B]; and Consejo Nacional de Ciencia Tecnologia e Innovacion tecnológica (CONCYTEC) [grant number 196-2013-FONDECYT/CONCYTEC] to [D.G.]. Funding for open access charge: Consejo Nacional de Ciencia Tecnologia e Innovacion tecnológica; Coordenação de Aperfeiçoamento de Pessoal de Nível Superior-Brasil; National Institutes of Health; and US Department of Energy Office of Basic Energy Sciences Nanomachine Program.

