## SUPPLEMENTARY MATERIALS, METHODS and DISCUSSION for "Interactions of Upstream and Downstream Promoter Regions with RNA Polymerase are Energetically Coupled and a Target of Regulation in Transcription Initiation"

**1. Open complexes form at the λ$p_R$ promoter and is active.** The wild type λ$p_R$ promoter contains an overlapping weaker secondary λ$p_{RM}$ promoter that starts transcription in the opposite direction that could compete with λ$p_R$ for the RNAP. However, AFM data showed that simultaneous binding on both promoters is a rare event, representing < 5% of DNA-protein complexes. Moreover, when it happens, it abolishes wrapping due to steric hindrance between the two polymerases (1). Consistently, the time-dependent production of specific RNA transcripts indicated that, in our experimental conditions, the majority of open complexes are active and formed at λ$p_R$ promoter. Transcription from λ$p_{RM}$ was undetectable (Figure S1A).

**2. Open complexes in AFM images.** AFM images displayed open complexes and non-specifc RNAP-DNA complexes (Figure S1B) even after incubation with heparin. We distinguished specific open complexes and non-specific RNAP-DNA complexes by the lengths of DNA at each side of the RNAP (arms) (2). Our sequence predicts a short-to-long DNA arm ratio of ~0.93 when the RNAP is bound at the λ$p_R$ promoter site in the 2 kbp-long DNA. However, in analyzing specific open complexes, we accepted arm ratios within ± 0.13 of the expected value (2). We found ~80% of complexes containing a single RNAP, and ~20% with two RNAPs. Open complexes with a single RNAP showed a DNA compaction of ~26 nm relative to free DNA (Figure S2A), consistent with previous reports for λ$p_R$ (2,3). Also, we detected ~50% of non-specific complexes that displayed similar or larger DNA compaction than those of specific open complexes. These complexes were identified by their random location along the 2 kbp DNA and discarded from the analysis (Figure S2B). It has been reported that non-specific RNAP-DNA complexes do not display DNA compaction (3). However, this discrepancy may reflect the different DNA sequences used in the experiments.

**3. Persistent length of DNA.** We determined the persistence length (P) of the 2 kbp-long DNA under our experimental conditions (20 mM Tris–HCl pH 7.9, 40 mM KCl, 5 mM $MgCl_2$, 1 mM DTT, 26$^oC$), using the WLC model (4,5). We obtained a value of P = 20.0 ± 5.6 nm, which is smaller than the 50 nm, often quoted in the literature for buffers that do not contain magnesium ions (6,7). Nonetheless, we found that for a short DNA fragment such as the one involved in the unwrapping/wrapping transitions, our estimation of the change in DNA contour length mean value, $\langle \Delta L_{rip} \rangle$, for P = 10, 20, 30, 40, and 50 nm differs at most by 1.3 nm (Figure S3).

**4. Characterization of RNAP-DNA complexes.** The unexpected detection of types for force-extension trace, displaying one and two transitions, hints at two different types of DNA-RNAP interactions (Figure S5A). Intriguingly, these two types of open complexes were not interconvertible, i.e. a complex with single transitions never displayed two transitions and viceversa. One scenario that could lead to the observation of two transitions is that the fragment contained one specific and one non-specific RNAP-DNA complexes, since we assembled open complexes with a 5-fold excess of RNAP over λ$p_R$. We performed several control experiments to discriminate and characterize specific open complexes at λ$p_R$ and non-specific RNAP-DNA complexes as follows.

- ✓ **4.1 Control I: Single molecule transitions correspond to RNAP-DNA complexes.** First, to determine if the observed transitions were due to RNAP binding to the template, we performed pulling/relaxing experiments with bare DNA in the absence of RNAP. No transitions were observed in this condition. Second, we performed pulling/relaxing experiments of a DNA mixed with RNAP previously incubated with heparin, which is known as a strong inhibitor of RNAP binding to DNA (8–10). Similarly, we observed no transitions and molecules instead displayed the force vs. extension behavior characteristic of bare DNA (4,11).
- ✓ **4.2 Control II: Low-force transtions correspond to non-specific complexes.** Next, we removed the λ$p_R$ promoter from the original 2 kbp-long DNA and incubated it with *E. coli*'s RNAP followed by the addition of heparin. We only observed molecules with a single transition (n = 180) in 2% of the total number of tethers formed (from 395 beads tested) with a transition force at 7.7 ± 1.6 pN and with evident hysteresis. These transitions survived for several pulling/relaxing cycles (> 30), and displayed changes in DNA extension ($\Delta x$) centered either at 25 or 35 nm (Figure S4). Coincidentally, these features ($\Delta x$, transition force, high hysteresis) are similar to the low-force

transitions present in curves that displayed two transitions under the standard conditions (see text and 1/2 in Figure S5A). These observations strongly suggest that the low force transitions correspond to non-specific RNAP-DNA interactions which are not inhibited by heparin. This finding is consistent with the fact that non-specific complexes also display DNA compaction in AFM images in our DNA template (Figure S2A).

- ✓ **4.3 Control III: Non-specific complexes display a different signature in a different DNA context.** The results from control II suggest that non-specific interactions display similar low force transition (< 8 pN) of force-extension traces displaying two transitions in Figure S5A. Therefore, we expect that in a different DNA context, non-specific interactions will vary accordingly, whereas the specific interactions with the λ*p*R promoter, should remain the same. To test this hypothesis, we repeated pulling/relaxing experiments of open complexes assembled on a similar 2 kbp-long DNA template where λ*p*R promoter remains at its center but flanked by different DNA sequences (context 2). The optical tweezers experiments revealed molecules with transition in 25% of the total tethers (Table S1) displaying unwrapping force at ~9 pN, with no significant hysteresis, $\Delta x$ = 15.5 nm, $\Delta L_U$ = 17.5 nm, and $\Delta G^0_{w\to u}$ = 13.7 kcal/mol (Figures S5 and S6, Tables S2-S5). These features are very similar to the high force transition of open complexes assembled in the original 2 kbp-long DNA (context 1). However, 25% of these molecules showed an additional non-cooperative low-force transition (Figure S5B). This result indicates that the non-specific RNAP-DNA interactions are DNA sequence-dependent and that the resulting non-specific complexes display transitions at a force below 8 pN either in a cooperative or non-cooperative manner. Therefore, we identify the high-force transition as a signature of the specific open complexes
- ✓ **4.4 Control IV: Specific open complexes are characterized by transitions at high-force.** Our results from control III strongly suggest that the high-force transition (Figures S5A-B) corresponds to specific binding of the RNAP to λ*p*R. To reinforce this observation, we redesigned our molecular construct to favor a 1:1 RNAP-DNA stoichiometry at λ*p*R. To this end, we assembled open complexes on a short DNA (231 bp-long DNA) bearing a single λ*p*R promoter extended from -170 to +61 bp. We then ligated the open complexes to two flanking DNA sequences (1 kbp-long each) labeled with digoxigenin and biotin, respectively (ligation in Figure S5C). When the ligation product was subjected to pulling/relaxing protocols, only 14% of total tethers showed a transition (n = 500). This transition showed no hysteresis at the force of ~9.6 pN, with $\Delta x$ ~15 nm, $\Delta L_U$ of 17.5 nm, and $\Delta G^0_{w\to u}$ of 13.4 kcal/mol (Figure S6, Tables S2-S5). These features are very similar to the high-force transition (Figure S5A-B) observed for open complexes assembled in either context 1 or 2. Thus, we confirm that specific transitions of open complexes are detected at forces of ~9 pN at standard salt conditions.

**5. The effect of ppGpp and glutamate on non-specific RNAP-DNA interactions.** The transcriptional regulator ppGpp is a small molecule that binds to RNAP and regulates gene expression by modulating the open complexes formation (2,12). Therefore, non-specific complexes should not be affected by the presence of ppGpp. As expected, ppGpp did not show a significant effect on $\Delta G^0_{w\to u}$ of non-specific RNAP-DNA complexes (Figure S4D). On the contrary, glutamate is a physiological anion that stabilizes both specific or non-specific DNA–protein interactions (13) and, as expected, glutamate increases the $\Delta G^0_{w\to u}$ value of these non-specific RNAP-DNA complexes (Figures S5A and S4D).

**6. Promoter occupancy of open complexes.** As stated above, we formed open complexes with a 5-fold excess of RNAP, therefore we expected high efficiency of finding open complexes in single molecule experiments. However, our yield was ~25% at low salt concentration (Table S1). This low efficiency can be due to: i) RNAP binding to the microfluidic chamber, and/or ii) RNAP dissociating from the promoter over time — open complexes lifetime is ~6 h (12). This interpretation is consistent with the fact that under more stabilizing conditions (e.g. in the presence of potassium glutamate) (9), our detection efficiency increased to ~45%; whereas under destabilizing conditions (e.g. λ$p_R$ mutants lacking UP elements (1)), our detection efficiency decreased to ~11%.

**7. Effective RNAP-λ$p_R$ association and transcription DNA bubble formation rates.** In our fluorescence assay, the increase of Cy3-fluorescence is due by RNAP-λ$p_R$ association (RNAP + λpR ↔ RPc) and DNA bubble formation ($I_{1L}$ ↔ $I_2$) (14), present in the $I_2$ and open complex. In the sequence of states leading to the formation of open complexes (RNAP + λpR ↔ RPc ↔ $I_{1E}$ ↔ $I_{1L}$ ↔ $I_2$ → RPo), all the intermediates are at equilibrium in the long time-frame and the resulting kinetics cannot be interpreted in terms of rate constants but as rates of equilibration. However, it is reasonable to assume that the states involved in the DNA strand opening ($I_{1L}$ → $I_2$) are not in equilibrium in the time-frame of the fluorescence experiment, because of the large energy barrier separating these two states, which makes that transition $I_{1L}$ → $I_2$ the rate-limiting step of the reaction RNAP + λpR → RPo. Under these conditions, it is appropriate to interpret the DNA bubble formation in term of as the forward rate ($k_o$) of DNA bubble formation: $I_{1L} \overset{k_o}{\rightarrow} I_2$. Additionally, since the late step of open complex formation ($I_2$ → RPo) is a fast process, $k_o$ also represents the rate of open complex formation ($I_{1L} \overset{k_o}{\rightarrow}$ RPo). Consider the simplified process of open complex formation,

$$D \overset{k_a}{\rightarrow} C \overset{k_o}{\rightarrow} O$$

being D the DNA (λ$p_R$), C the closed complex and O the open complex. Here $k_a$ is an effective RNAP-λ$p_R$ association rate constant since it is a fast and reversible process,

$$k_a = \frac{k_1[RNAP]k_o}{k_2 + k_o}$$

Where $k_1$ and $k_2$ are the RNAP-λ$p_R$ association and dissociation rate constants, respectively. [RNAP] is the RNAP concentration.

The kinetic equations for λ$p_R$ and RNAP binding is defined by,

$$D(t) = D_0 exp\{-k_a t\}$$

for the closed complex formation,

$$C(t) = \frac{k_a D_0}{k_o - k_a}\{exp(-k_a t) - exp(-k_o t)\}$$

and for the open complex formation,

$$O(t) = D_0 \left\{1 - \frac{1}{k_o - k_a}[k_o exp(-k_a t) + k_a exp(-k_o t)]\right\}$$

Experimentally, we observe that Cy3-fluorescence increases by a certain factor $\beta$ when RNAP is bound and by a factor $\alpha$ when DNA bubble is opened. In the absence of RNAP, the basal Cy3 fluorescence is proportional to λ$p_R$ concentration by a factor of $\delta$. Therefore, the total fluorescence in time would be,

$$F(t) = \delta D(t) + \beta C(t) + \alpha O(t)$$

Substituting previous equations, we have

$$F(t) = \alpha D_0 + \left\{\delta D_0 - \alpha \frac{D_0 k_o}{k_o - k_a} + \beta \frac{k_a D_0}{k_o - k_a}\right\} exp\{-k_a t\} + \left\{\alpha \frac{D_0 k_a}{k_o - k_a} - \beta \frac{k_a D_0}{k_o - k_a}\right\} exp(-k_o t)$$

recalling,

$$\delta D_0 - \alpha \frac{D_0 k_o}{k_o - k_a} + \beta \frac{k_a D_0}{k_o - k_a} = A$$

$$\alpha \frac{D_0 k_a}{k_o - k_a} - \beta \frac{k_a D_0}{k_o - k_a} = B$$

$$\alpha D_0 = C$$

we obtain,

$$F(t) = C + Aexp\{-k_a t\} + Bexp(-k_o t)$$

Which is similar to the empirical fitting equation reported in the literature (14). Note that $k_a$ and $k_o$ are effective rates constant of RNAP-λ$p_R$ association and DNA bubble formation, respectively.

### SUPPLEMENTARY FIGURES DESCRIPTION

**Figure S1: Open complex transcription activity and representative Atomic Force Microscopy (AFM) micrographs of open complexes. (A)** Activity assay of open complexes in the 231 bp-long DNA, bearing the λ$p_R$ promoter. When RNA polymerase (RNAP) forms an open complex at λ$p_R$, it produces a 91nt RNA transcript (top red arrow), and a 56nt transcript (bottom red arrow) when it transcribes from the λ$p_{RM}$ promoter. As expected, most of the RNA transcript derives from λ$p_R$. Negative control: when RNAP is previously incubated with heparin, we do not observe RNA transcript, indicating that open complex was not formed (**B)** AFM image of open complexes formed with λ$p_R$ promoter and *E. coli*'s RNAP-$\sigma^{70}$ in the 2 kbp DNA. (**C)** Gallery of four open complexes, the RNAP is seen near the center of the template where λ$p_R$ promoter is located. Scale bars are shown for each picture. White arrows indicate RNAP and DNA.

**Figure S2: AFM controls. (A)** Distributions of the contour length of bare DNA (top panel, N = 504 molecules), specific (medium panel, N = 153), and non-specific complexes (bottom panel, N = 150) measured from AFM images. Red arrow indicates that there is a second population of non-specific complexes with a shorter contour length. Black line is the Gaussian fitting. We observed a DNA compaction of 26.3 ± 2.5 nm in open complex relative to free DNA. **(B)** short-to-long DNA arm ratio distributions of all RNAP-DNA complexes (top panel), open (medium panel) and non-specific RNAP-DNA complexes (bottom panel). Specific open complex has a mean ratio of 0.9 ± 0.01 (SD = 0.1, N = 153).

**Figure S3: Effect of the persistent length on contour length estimation.** Histograms of the change in DNA contour length, $\Delta L_{rip}$, derived from transitions of open complexes formed at λ$p_R$ promoter at standard salt conditions (40mM KCl). The $\Delta L_U$ was obtained by fitting the change in DNA extension ($\Delta x$) at the observed transition force ($F_U$) using the WLC model, with persistent lengths (P) of 10, 20, 30, 40 and 50 nm. The resulting $\langle \Delta L_U \rangle$ values are indicated. Note that $\Delta L_U$ does not change significantly in this range of P values.

**Figure S4: Characterization of force-extension curves of non-specific RNAP-DNA complexes. (A)** Pulling (blue) and relaxing (red) cycles of promoter-less 2 kbp DNA incubated with RNAP and subsequently incubated with heparin. Red arrows indicate the expected position of the high force transition observed in specific open complexes (Fig. S5). Black arrows indicate the positions of low force transitions of non-specific complexes (Fig. S5). All RNAP-DNA complexes prepared with promoter-less DNA as shown in this figure displayed no transition or a single transition. Only 4% of total tethers displayed transitions (N = 180). **(B)** Distribution of change in DNA extension ($\Delta x$) of non-specific complexes, showing two peaks, one with a mean of 24.3 nm ($\sigma_{\Delta x}$ = 5.4 nm) and the other with a mean of 35.1 nm ($\sigma_{\Delta x}$ = 6.0 nm). **(C)** Transition force distribution of non-specific complexes, showing a mean of a 7.7 pN ($\sigma_F$ = 1.6 pN). **(D)** Comparison of thermodynamic stability of non-specific complexes (Fig. S5A-B) in standard conditions (top panel) with their stability in conditions known to stabilize (100mM of glutamate) or destabilize (100uM ppGpp) open complexes. Work distributions for disrupting (red) and forming (blue) DNA-RNAP contacts. Histograms were equally binned for each condition; points and lines refer to experimental values and Gaussian fittings, respectively. Note that, while glutamate stabilizes non-specific DNA-RNAP complexes, ppGpp does not have a significant effect on those, as expected.

**Figure S5. Comparison of transitions of open complexes formed on various DNA templates to probe open complex specificity. (A)** Force-extension pulling (blue) and relaxing (red) curves of open complexes formed in a 2 kbp-long DNA bearing the λ$p_R$ promoter in the middle (Context 1). **(B)** A different 2 kbp DNA template where the λ$p_R$ promoter is located also in the middle but its flanking DNA sequence is varied (Context 2). **(C)** The open complex is also obtained on a 231 bp-long DNA bearing the λ$p_R$ promoter and then ligated to two 1 kbp DNA flanking sequences (Ligation). In the latter construct, only the high force transition was observed.

**Figure S6. Comparative analysis of open complexes formed in context 1 (top), context 2 (medium) and a ligation product (bottom panel). (A)** Change in DNA extension ($\Delta x$) distributions. **(B)** Transition force distributions during pulling experiments. **(C)** Work distributions of disrupting (red) and forming (blue) promoter-RNAP contacts. Black, red and blue curves are Gaussian fittings. For full description of statistics, see Table S2-S5.

**Figure S7: Characterization of open complexes presenting two transitions, the low and high force. (A)** Transition force ($F_U$, pN) vs change in DNA extension ($\Delta x$, nm) plots. $F_U$ and $\Delta x$ histogram distributions for the low and high force transition, are showed in cyan and purple, respectively. **(B)** Work distributions for disrupting (red) and forming (blue) promoter-RNAP contacts of the high (top panel) and low force transition (bottom panel). Histograms are equally binned for each condition; points and lines refer to experimental values and Gaussian fittings, respectively. For full description of statistics, see Table S5-S6.

**Figure S8: TEM micrographs of negatively stained of open complexes. (A)** Micrograph patch where two RPo are marked by red circles. **(B), (C)** Detail of circled RPo in a. Scale bar is shown.

**Figure S9**: **3D reconstruction and model of RPo. (A)** EM reconstruction of the open complex (0.42 σ contour level) indicating the β dispensable region 1 (βDR1), β dispensable region 2 (βDR2), β' insert 6 (β'i6), $\sigma^{70}$ non-conserved region ($\sigma^{70}$NCR), αNTD dimer, and the ω subunit. We use the "hide-dust" tool of Chimera software to remove those densities not connected to the EM model at 0.42 σ contour level. **(B)** Fourier Shell Correlation (FSC) with 1/2-bit and 0.143 criterion indicating the resolution of the EM model at 17 Å and 16 Å, respectively. **(C)** The open complex EM map with a contour level at 6 σ (purple semi-transparent surface), indicating the wrapping path located upstream from the transcription bubble. The DNA promoter and RNAP coordinate models are shown in black and light-gray ribbon respectively, and are fitted in the EM map at 0.42 σ contour level (gray transparent mesh). Densities not related to the DNA path are not shown. The observation of this strong density, even in a high contour level EM map, is probable due to the high affinity of the uranyl acetate for DNA as reported in another study (15).

**Figure S10: Selected 2D class-averages of the open complex. (A)** Representative 2D class-averages (left), the corresponding 2D re-projections (center) and the fitted coordinate model in the same orientation (right). The labels correspond to domains and subunits in the 3D map. The color code is as follows: β subunit (magenta ribbon), β' subunit (green ribbon), the $\alpha^{I}$NTD (blue ribbon), $\alpha^{II}$NTD (cyan ribbon), $\sigma^{70}$ (yellow ribbon) and the ω subunit (red ribbon). The overall DNA promoter is shown in orange ribbon. **(B)** Other class-averages (top row) and their respective re-projections (bottom row) used in the final 3D structure reconstruction.

**Figure S11: Additional pulling/relaxing trajectories of two open complexes.** Five examples of consecutive force-extension pulling (blue) and relaxing (red) curves of two different open complexes (A and B) formed in a 2 kbp-long DNA bearing the λ$p_R$ promoter wild-type in the middle.

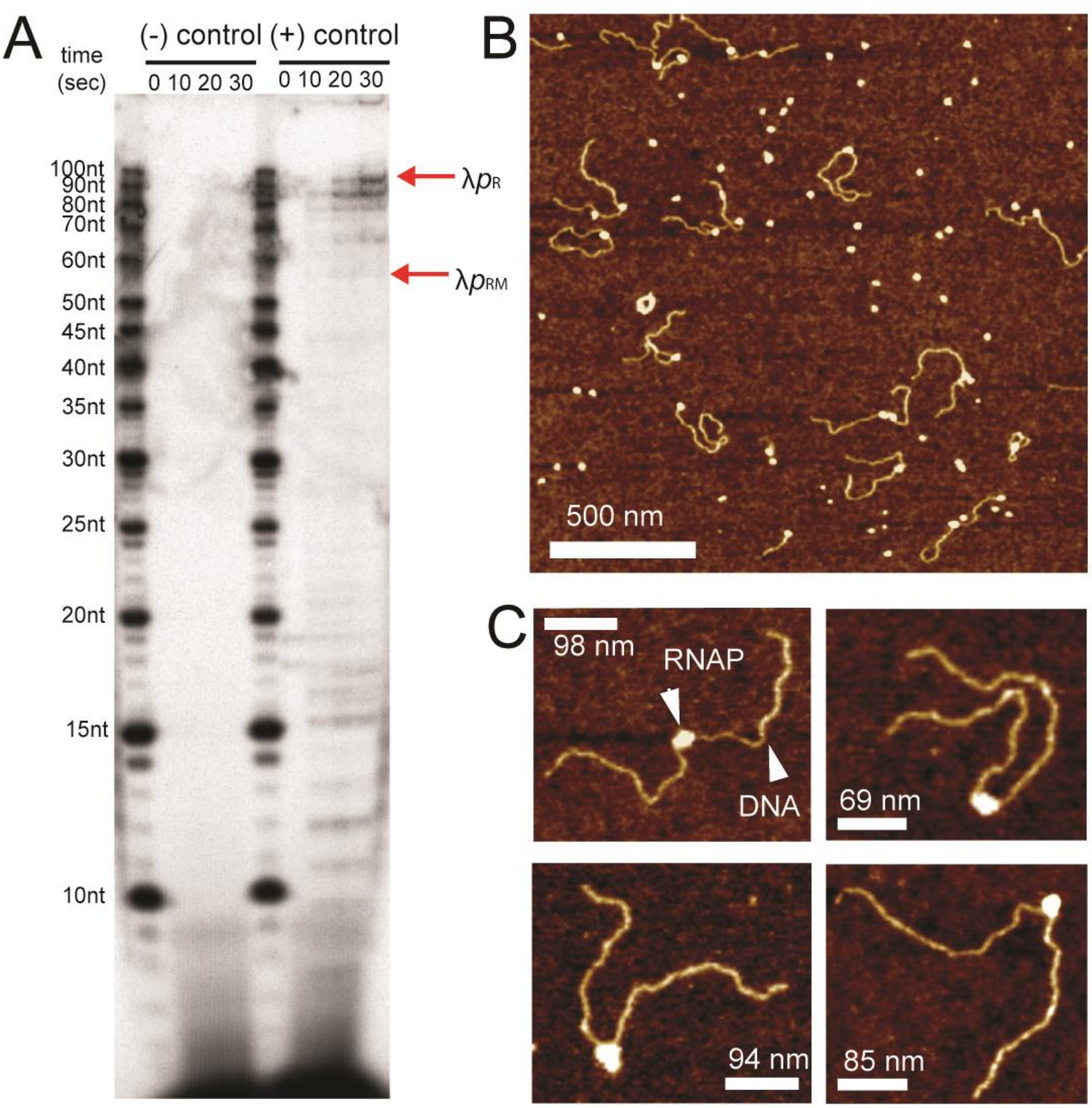
A
time (sec)
(-) control
(+) control
0 10 20 30
0 10 20 30
100nt
90nt
80nt
70nt
60nt
50nt
45nt
40nt
35nt
30nt
25nt
20nt
15nt
10nt
λpR
λpRM
B
500 nm
C
98 nm
RNAP
DNA
69 nm
94 nm
85 nm

**Figure S1**

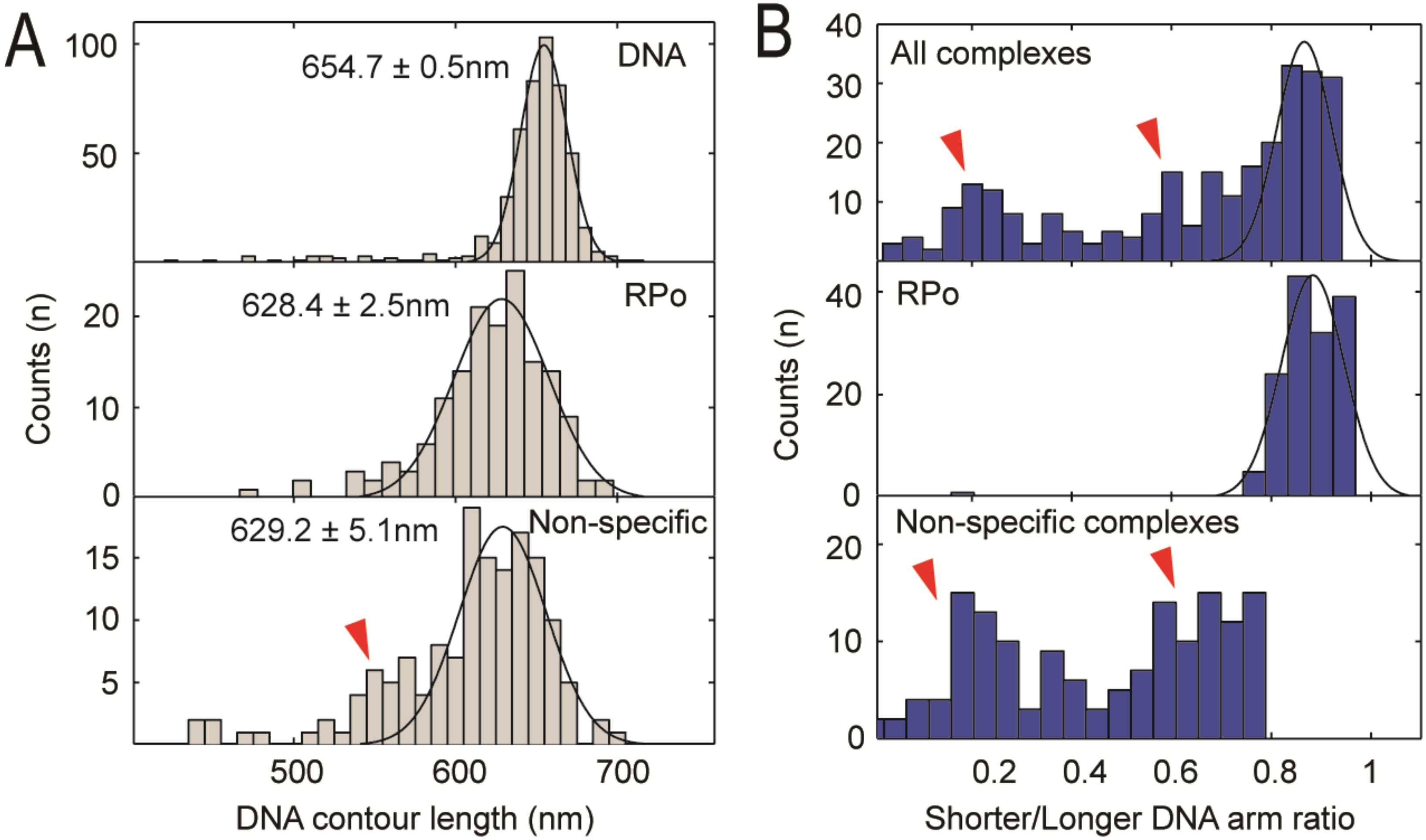
A
B
Counts (n)
654.7 ± 0.5nm
DNA
628.4 ± 2.5nm
RPo
629.2 ± 5.1nm
Non-specific
DNA contour length (nm)
All complexes
RPo
Non-specific complexes
Shorter/Longer DNA arm ratio

**Figure S2**

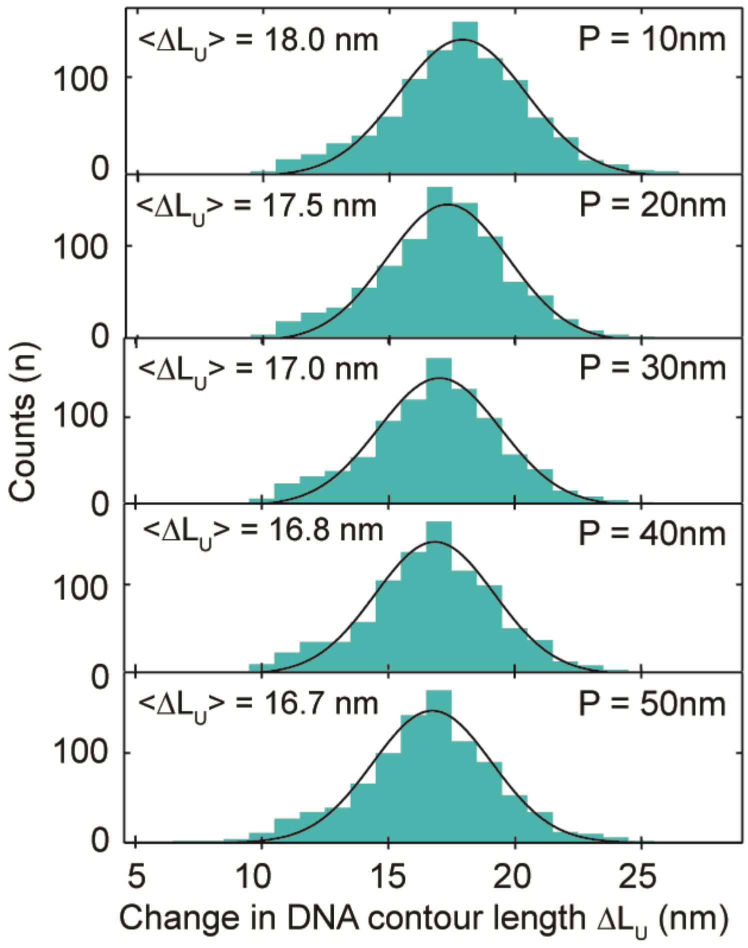

<ΔL_U> = 18.0 nm
P = 10nm
<ΔL_U> = 17.5 nm
P = 20nm
<ΔL_U> = 17.0 nm
P = 30nm
<ΔL_U> = 16.8 nm
P = 40nm
<ΔL_U> = 16.7 nm
P = 50nm
Counts (n)
100
0
5
10
15
20
25
Change in DNA contour length ΔL_U (nm)


**Figure S3**

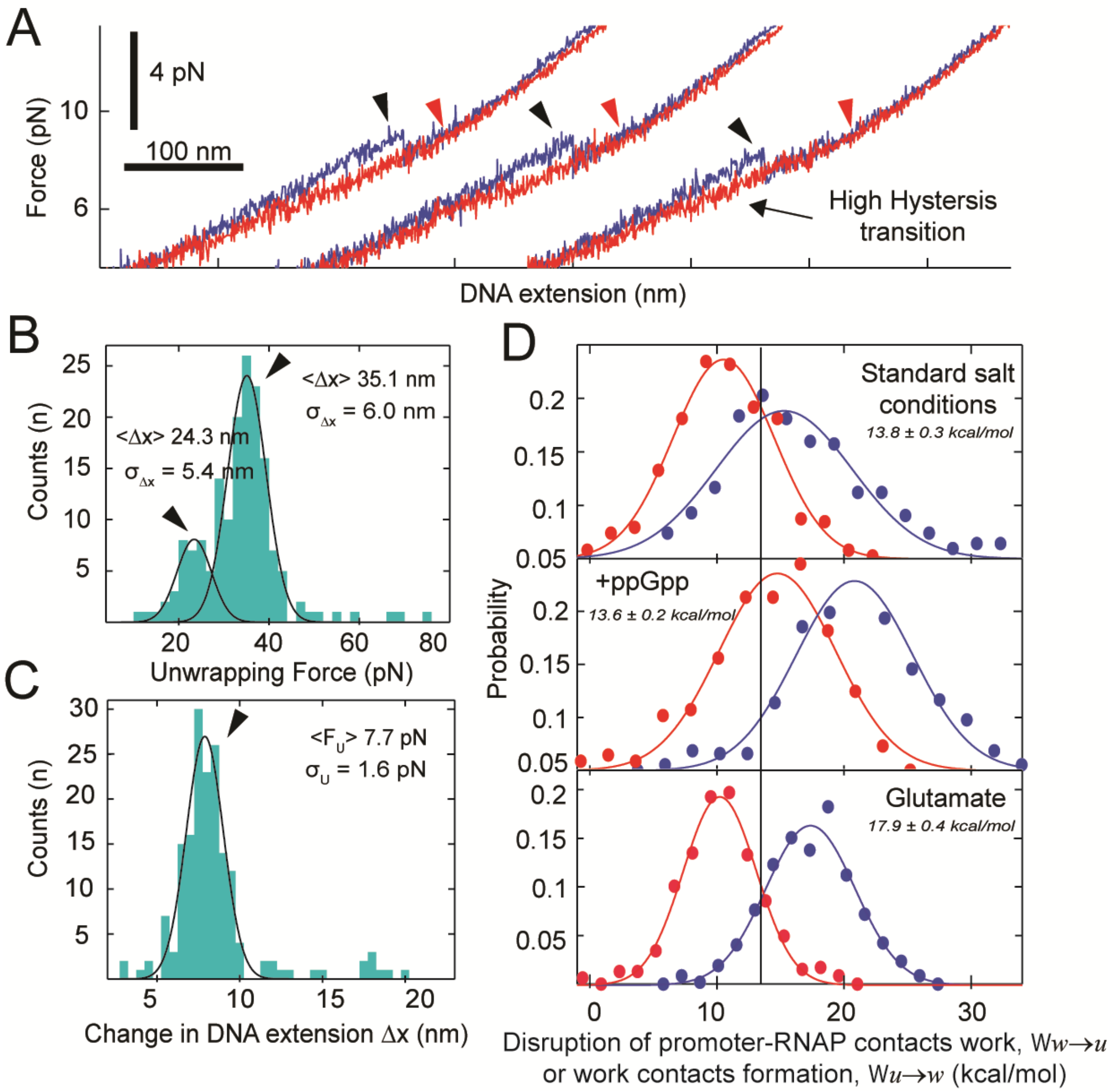
A
4 pN
100 nm
Force (pN)
10
6
High Hysteresis transition
DNA extension (nm)
B
Counts (n)
25
20
15
10
5
<Δx> 35.1 nm
σ_Δx = 6.0 nm
<Δx> 24.3 nm
σ_Δx = 5.4 nm
20
40
60
80
Unwrapping Force (pN)
C
Counts (n)
30
25
20
15
10
5
<F_U> 7.7 pN
σ_U = 1.6 pN
5
10
15
20
Change in DNA extension Δx (nm)
D
Standard salt conditions
13.8 ± 0.3 kcal/mol
+ppGpp
13.6 ± 0.2 kcal/mol
Glutamate
17.9 ± 0.4 kcal/mol
Probability
0.2
0.15
0.1
0.05
0
10
20
30
Disruption of promoter-RNAP contacts work, Ww→u
or work contacts formation, Wu→w (kcal/mol)

**Figure S4**

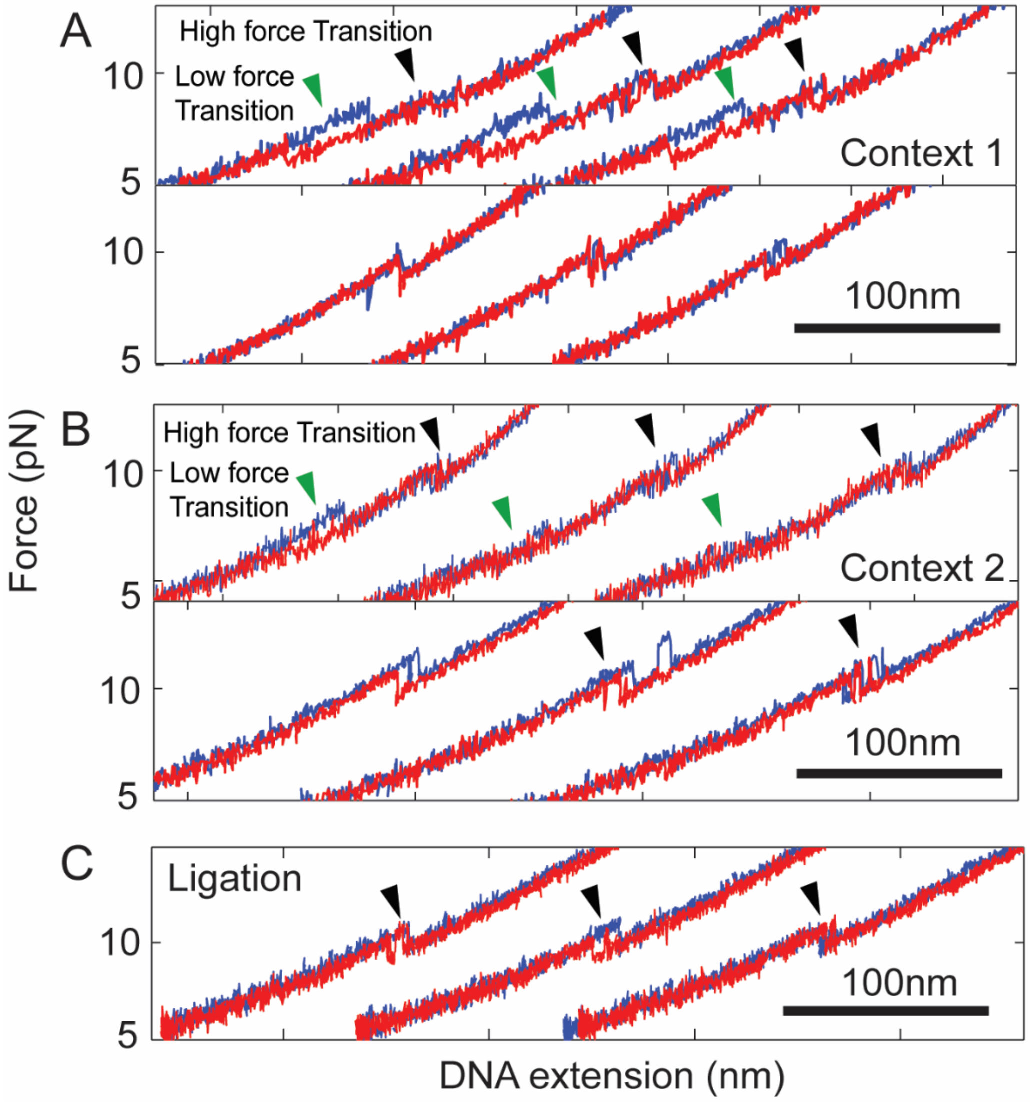
A
High force Transition
Low force
Transition
Context 1
100nm
B
High force Transition
Low force
Transition
Context 2
100nm
C
Ligation
100nm
Force (pN)
DNA extension (nm)
10
5

**Figure S5**

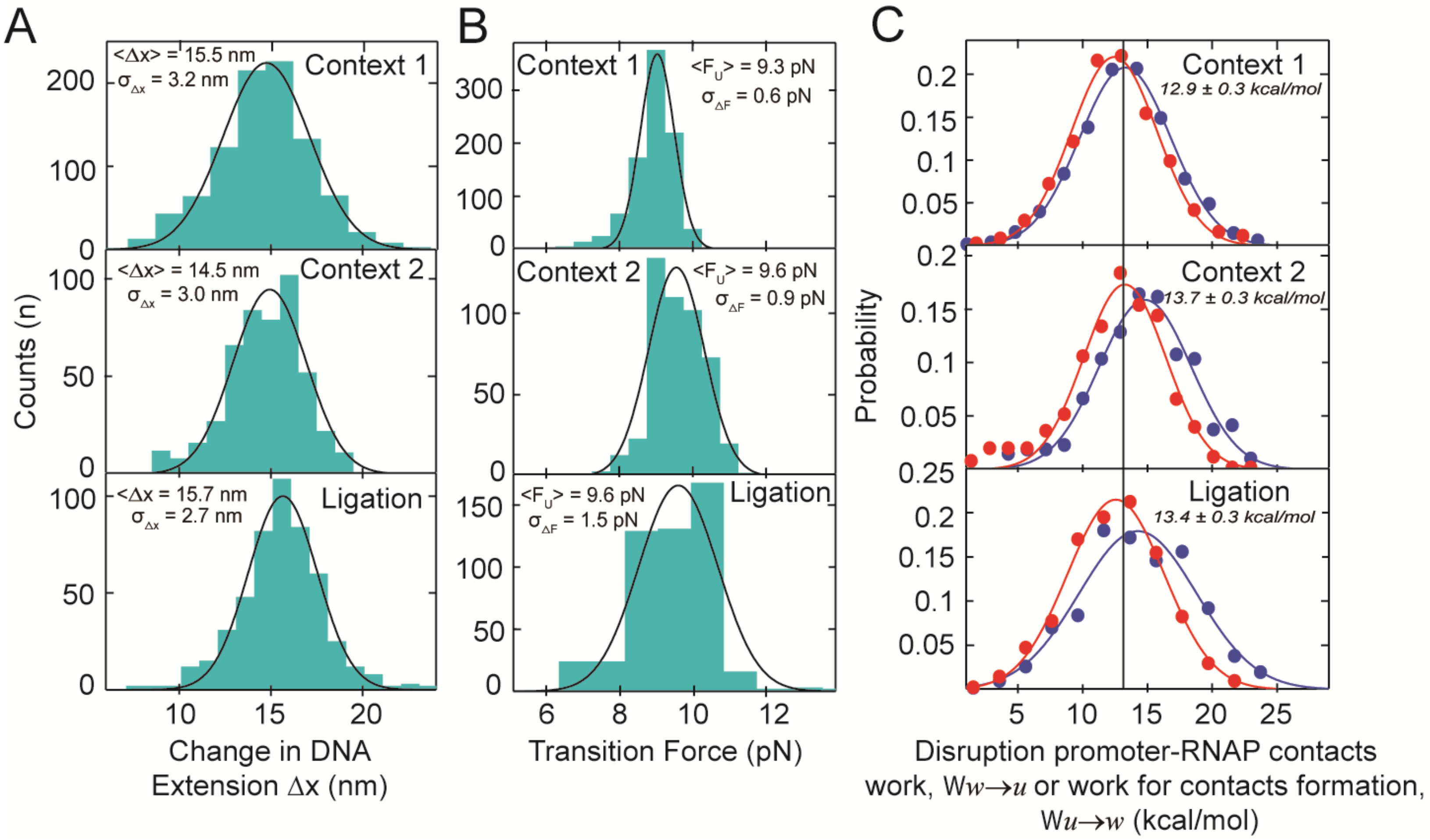
A
B
C
Counts (n)
<Δx> = 15.5 nm
σΔx = 3.2 nm
Context 1
<Δx> = 14.5 nm
σΔx = 3.0 nm
Context 2
<Δx = 15.7 nm
σΔx = 2.7 nm
Ligation
Change in DNA
Extension Δx (nm)
Context 1
<FU> = 9.3 pN
σΔF = 0.6 pN
Context 2
<FU> = 9.6 pN
σΔF = 0.9 pN
<FU> = 9.6 pN
σΔF = 1.5 pN
Ligation
Transition Force (pN)
Probability
Context 1
12.9 ± 0.3 kcal/mol
Context 2
13.7 ± 0.3 kcal/mol
Ligation
13.4 ± 0.3 kcal/mol
Disruption promoter-RNAP contacts
work, Ww→u or work for contacts formation,
Wu→w (kcal/mol)

**Figure S6**

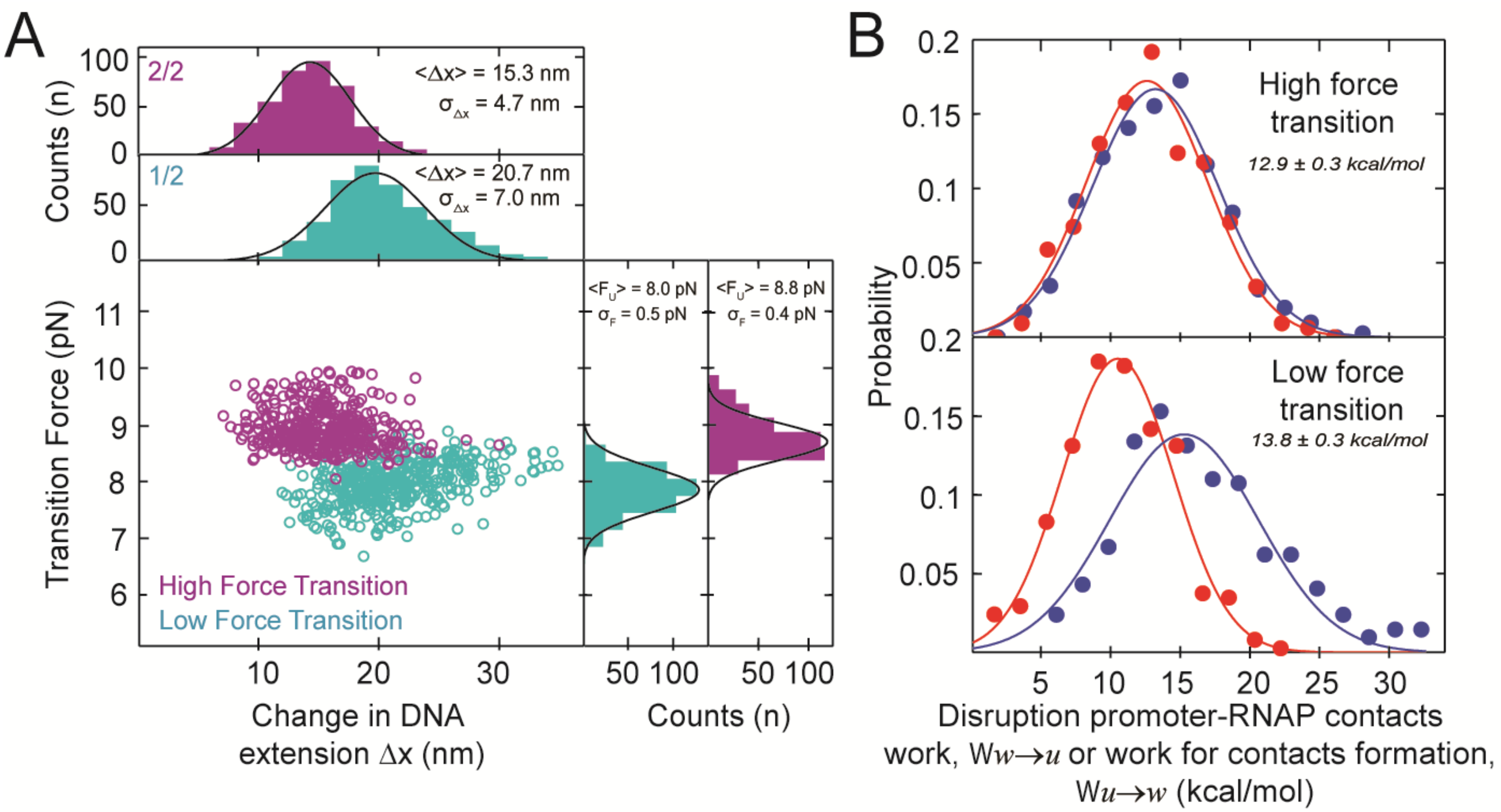

A
B
Counts (n)
2/2
<Δx> = 15.3 nm
σ Δx = 4.7 nm
1/2
<Δx> = 20.7 nm
σ Δx = 7.0 nm
Transition Force (pN)
<F U> = 8.0 pN
σ F = 0.5 pN
<F U> = 8.8 pN
σ F = 0.4 pN
High Force Transition
Low Force Transition
Change in DNA extension Δx (nm)
Counts (n)
Probability
High force transition
12.9 ± 0.3 kcal/mol
Low force transition
13.8 ± 0.3 kcal/mol
Disruption promoter-RNAP contacts work, Ww→u or work for contacts formation, Wu→w (kcal/mol)


**Figure S7**

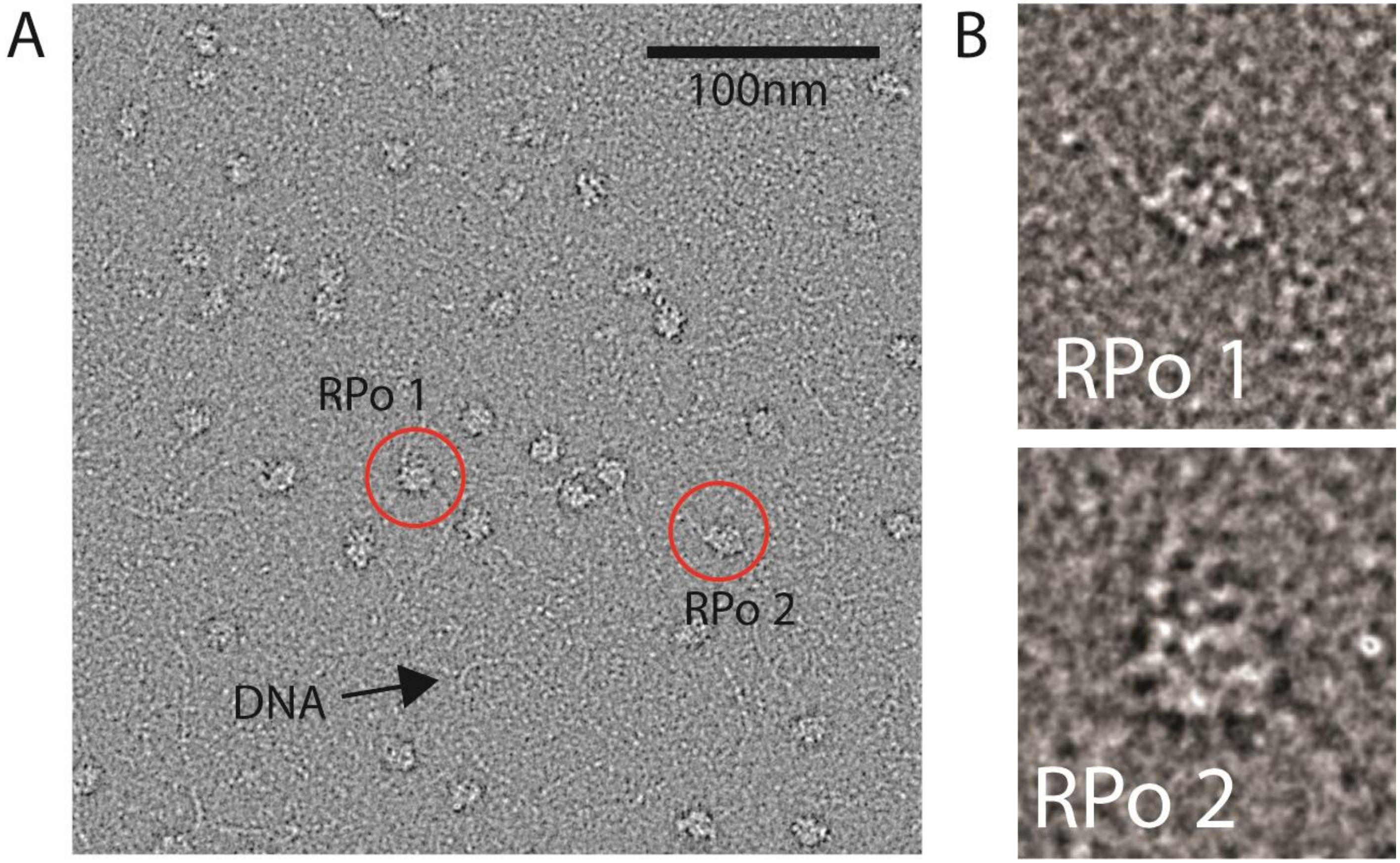


**Figure S8**

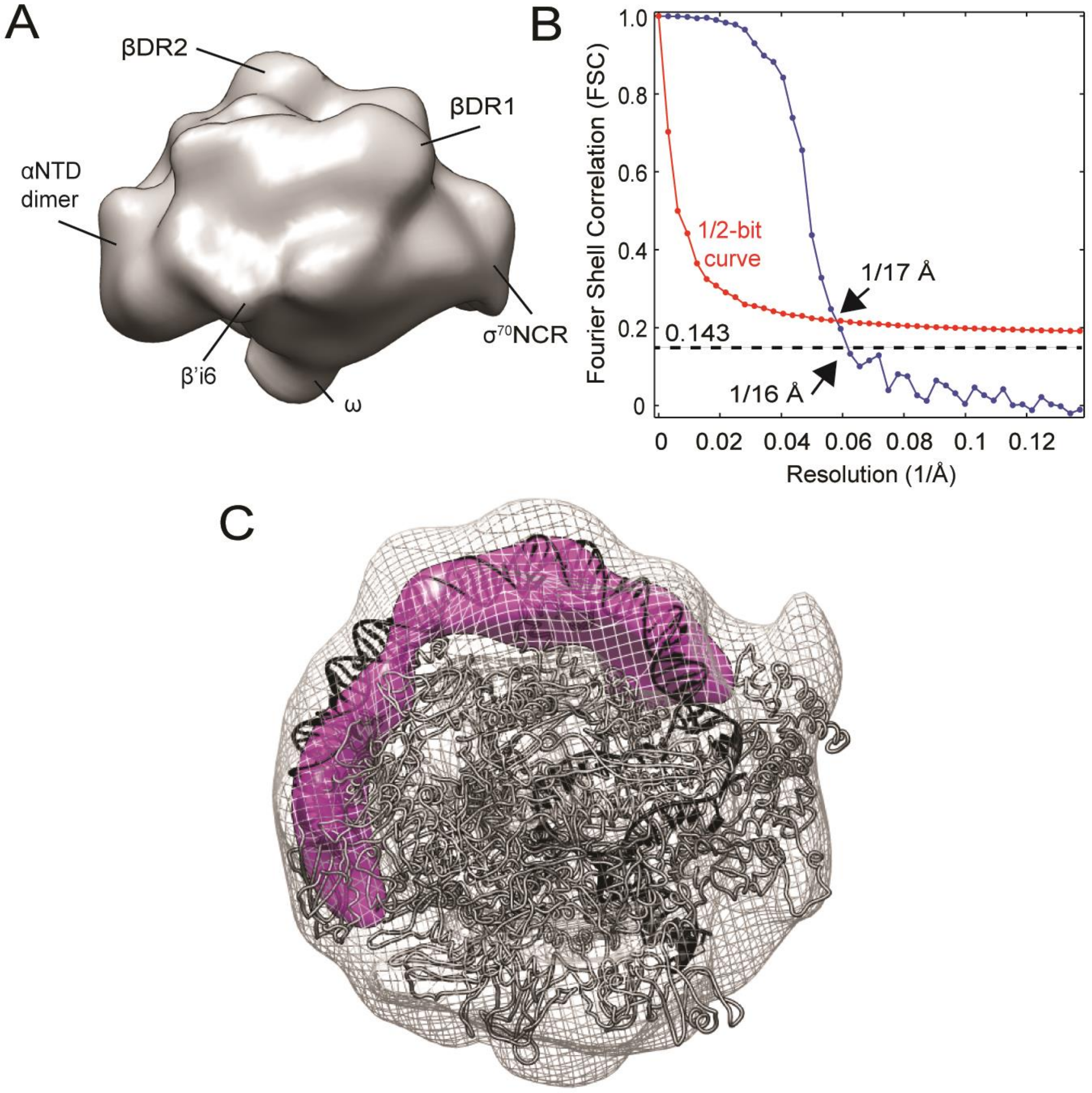
A
βDR2
βDR1
αNTD
dimer
$\sigma^{70}$NCR
β'i6
ω
B
Fourier Shell Correlation (FSC)
1.0
0.8
0.6
0.4
0.2
0
1/2-bit
curve
1/17 Å
0.143
1/16 Å
0
0.02
0.04
0.06
0.08
0.1
0.12
Resolution (1/Å)
C

**Figure S9**

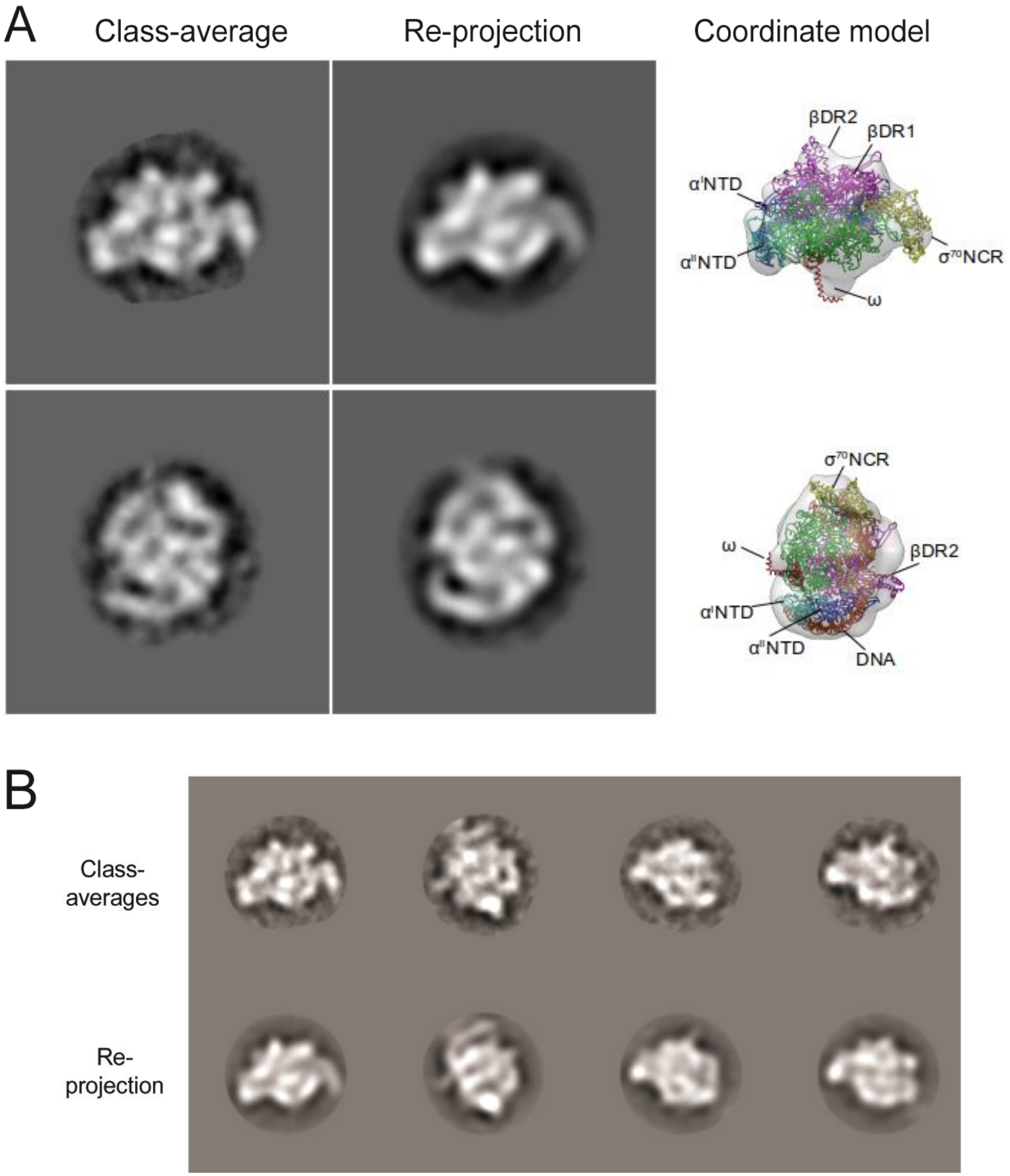
A
Class-average
Re-projection
Coordinate model
βDR2
βDR1
α^I NTD
α^II NTD
σ^70 NCR
ω
σ^70 NCR
ω
βDR2
α^I NTD
α^II NTD
DNA
B
Class-averages
Re-projection

**Figure S10**

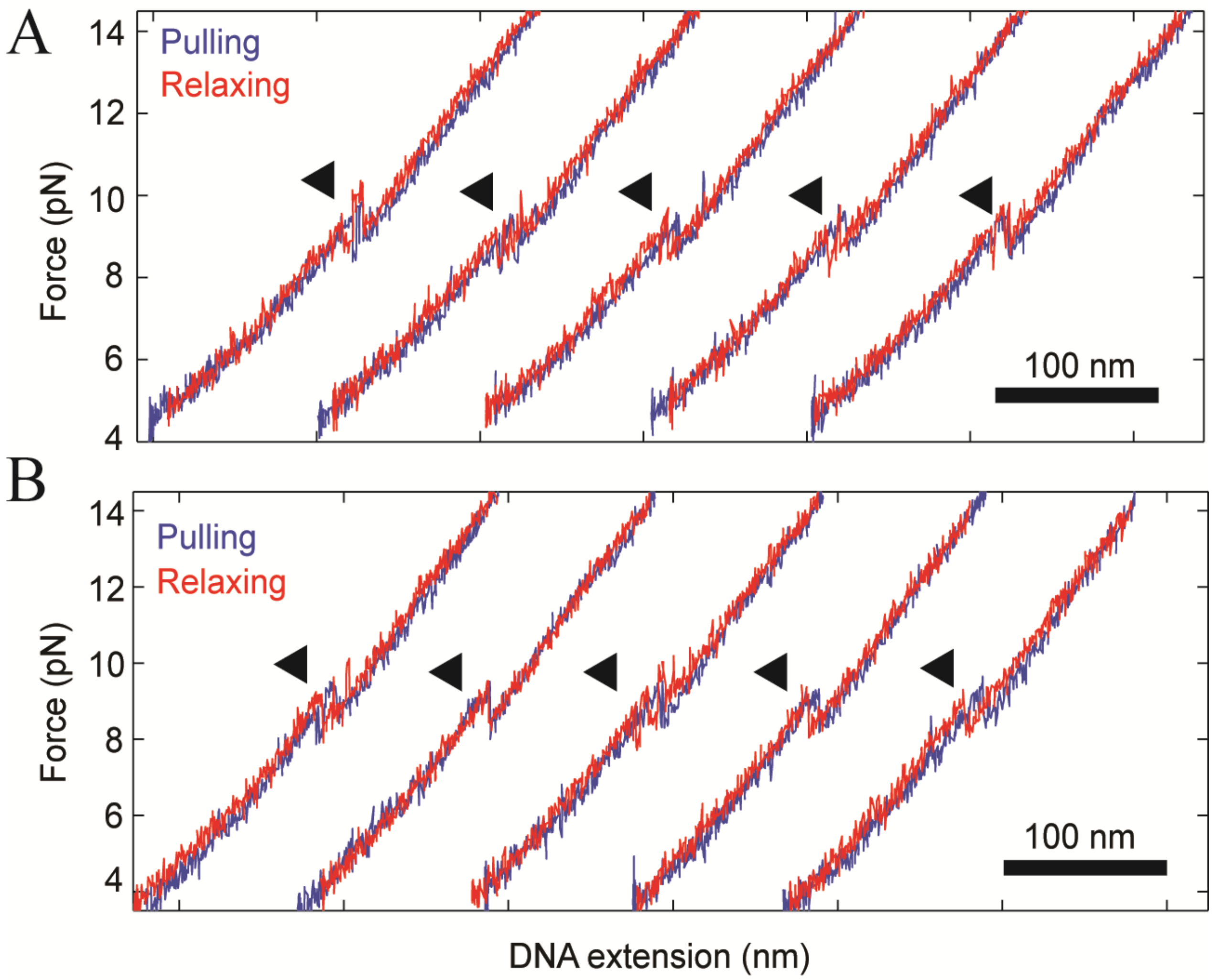
A
14
12
10
8
6
4
Force (pN)
Pulling
Relaxing
100 nm
B
14
12
10
8
6
4
Force (pN)
Pulling
Relaxing
100 nm
DNA extension (nm)

**Figure S11**

**Table S1:** Efficient of finding open complexes in optical tweezers experiments

| Promoter | Condition | n* | N** | Percentage of tethers displaying open complexes*** |
|---|---|---|---|---|
| **λ$p_R$ wt (Context 1)** | 40mM KCl | 920 | 29 | 21% |
| **λ$p_R$ wt (Context 2)** | | 678 | 15 | 25% |
| **λ$p_R$ wt ligation (Context 1)** | | 500 | 14 | 14% |
| **λ$p_R$ mutA*** ** | | 371 | 12 | 17% |
| **λ$p_R$ mutB*** ** | | 226 | 6 | 11% |
| **λ$p_R$ wt*** ** | 100mM KGlu | 1182 | 25 | 43% |
| | 300mM KCl | 510 | 15 | 27% |
| | 100uM ppGpp | 597 | 12 | 20% |

* n = Total number of transitions analyzed.
** N = Total number of open complexes analyzed. All of them were single tether.
*** Percentage of open complexes per tether tested, they include double and single tether and also noisy data. The remaining tethers that did not displayed transitions behaved mechanically as naked DNA

**Table S2**: Summary of transition forces during the pulling/relaxing experiments

| Promoter | Condition | $\langle F_U \rangle$ (± e)* | $\sigma_{F,U}$ + | $R^2$(U)++ | $\langle F_W \rangle$ (± e)* | $\sigma_{F,W}$ + | $R^2(W)$++ | n◊ | N◊◊ |
|---|---|---|---|---|---|---|---|---|---|
| **λ$p_R$ wt (Context 1)** | 40mM KCl | 9.3 ± 0.1 | 0.6 | 0.96 | 9.0 ± 0.1 | 0.7 | 0.98 | 920 | 29 |
| **λ$p_R$ wt (Context 2)** | | 9.6 ± 0.1 | 0.9 | 0.99 | 9.2 ± 0.1 | 0.9 | 0.99 | 678 | 15 |
| **λ$p_R$ wt ligation (Context 1)** | | 9.6 ± 0.3 | 1.5 | 0.89 | 9.3 ± 0.1 | 1.0 | 0.97 | 500 | 14 |
| **λ$p_R$ mutA**** | | 11.1 ± 0.1 | 1.1 | 0.99 | 10.7 ± 0.1 | 1.0 | 0.99 | 371 | 12 |
| **λ$p_R$ mutB**** | | 10.5 ± 0.1 | 1.2 | 0.98 | 8.7 ± 0.1 | 1.1 | 0.96 | 226 | 6 |
| **λ$p_R$ wt**** | 100mM KGlu | 10.7 ± 0.1 | 0.9 | 0.94 | 10.2 ± 0.1 | 0.9 | 0.94 | 1182 | 25 |
| | 300mM KCl | 9.7 ± 0.1 | 0.7 | 0.99 | 9.5 ± 0.1 | 0.7 | 0.99 | 510 | 15 |
| | 100μM ppGpp | 10.0 ± 0.1 | 0.6 | 0.88 | 9.8 ± 0.1 | 0.7 | 0.89 | 597 | 17 |

* Mean transition forces (pN), $\langle F_U \rangle$ (± error) and $\langle F_W \rangle$ (± error) respectively.
** These experiments were performed in context 1.
+ Standard deviation of transition forces (pN) distribution, $\sigma_{F,U}$ and $\sigma_{F,W}$
++ $R^2$ from normal fitting to transition forces (pN) distribution.
◊ Number of transitions per condition
◊◊ Number of open complexes.

**Table S3:** Summary of change in DNA extension of transitions during the pulling/relaxing experiments

| Promoter | Condition | $\langle \Delta x \rangle_U$* | $\sigma_{\Delta x,U}$+ | $R^2(U)$++ | $\langle \Delta x \rangle_W$ | $\sigma_{\Delta x,W}$+ | $R^2(W)$++ | n◊ | N◊◊ |
|---|---|---|---|---|---|---|---|---|---|
| **λ*p*R wt (Context 1)** | 50mM KCl | 15.5 ± 0.2 | 3.2 | 0.98 | 15.1 ± 0.1 | 3.2 | 0.99 | 920 | 29 |
| **λ*p*R wt (Context 2)** | | 14.5 ± 0.2 | 3.0 | 0.99 | 14.1 ± 0.2 | 2.9 | 0.98 | 678 | 15 |
| **λ*p*R wt ligation (Context 1)** | | 15.7 ± 0.2 | 2.7 | 0.98 | 15.0 ± 0.1 | 2.5 | 0.99 | 500 | 14 |
| **λ*p*R mutA**** | | 16.8 ± 0.1 | 2.1 | 0.99 | 16.6 ± 0.1 | 1.8 | 0.98 | 371 | 12 |
| **λ*p*R mutB**** | | 11.4 ± 0.1 | 3.6 | 0.98 | 10.3 ± 0.3 | 2.9 | 0.98 | 226 | 6 |
| **λ*p*R wt**** | 100mM KGlu | 15.9 ± 0.3 | 3.5 | 0.98 | 15.6 ± 0.3 | 3.2 | 0.98 | 1182 | 25 |
| | 300mM KCl | 12.6 ± 0.1 | 2.5 | 0.99 | 12.2 ± 0.1 | 2.7 | 0.99 | 510 | 15 |
| | 100uM ppGpp | 11.4 ± 0.1 | 2.6 | 0.99 | 11.1 ± 0.2 | 2.6 | 0.99 | 597 | 12 |

* Mean of change in DNA extension of transitions (nm), $\langle \Delta x \rangle_U$ (± error) and $\langle \Delta x \rangle_W$ (± error), respectively.

** These experiments were performed in context 1.

+ Standard deviation of change in DNA extension of transitions (nm), $\sigma_{\Delta x,U}$ and $\sigma_{\Delta x,W}$

++ $R^2$ from normal fitting to the change in DNA extension of transitions (nm) distribution, $R^2(U)$ and $R^2(W)$, respectively.

◊ Number of transitions per condition.

◊◊ Number of open complexes.

**Table S4:** Summary of the Change in DNA contour length of transitions of open complexes

| Promoter | Condition | Disruption of promoter-RNAP interactions ($u \rightarrow w$) | | | Formation of promoter-RNAP interactions ($u \rightarrow w$) | | | n[++] | N[++] |
|---|---|---|---|---|---|---|---|---|---|
| | | $\langle \Delta L_U \rangle$* | $\sigma_{\Delta L_U}$** | P-value’ | $\langle \Delta L_W \rangle$* | $\sigma_{\Delta L_W}$** | P-value’ | | |
| **λ*p*$_R$ wt (context 1)** | 40mM KCl | 17.5 ± 0.3 | 3.7 | --- | 17.5 ± 0.2 | 3.6 | --- | 920 | 29 |
| **λ*p*$_R$ wt (context 2)** | | 17.0 ± 0.1 | 3.4 | 0.0 | 17.5 ± 0.2 | 3.6 | 0.0 | 678 | 15 |
| **λ*p*$_R$ wt (Ligation)** | | 17.5 ± 0.2 | 3.1 | 0.0 | 17.0 ± 0.2 | 3.2 | 0.0 | 500 | 14 |
| **λ*p*$_R$ mutA*** ** | | 17.5 ± 0.2 | 3.1 | 1.0 | 17.5 ± 0.3 | 2.8 | 0.3 | 500 | 14 |
| **λ*p*$_R$ mutB*** ** | | 13.5 ± 0.3 | 3.9 | 1.0 | 12.5 ± 0.1 | 3.0 | 1.0 | 226 | 6 |
| **λ*p*$_R$ wt*** ** | 100mM KGlu | 18.6 ± 0.3 | 3.4 | 1.0 | 18.6 ± 0.2 | 3.5 | 1.0 | 1182 | 25 |
| | 300mM KCl | 15.0 ± 0.1 | 2.7 | 1.0 | 14.0 ± 0.1 | 2.9 | 1.0 | 510 | 15 |
| | 100uM ppGpp | 13.7 ± 0.1 | 2.6 | 1.0 | 13.4 ± 0.1 | 2.8 | 1.0 | 597 | 12 |

***** Mean of change in DNA contour length of transitions, $\langle \Delta L_{rip} \rangle$ and $\langle \Delta L_{zip} \rangle$ (± error), respectively. It was estimated using the WLC model, $F = \frac{K_B T}{P}\left\{\frac{1}{4}\left(1-\frac{\Delta x}{\Delta L}\right)^{-2} - \frac{1}{4} + \frac{\Delta x}{\Delta L}\right\}$, where P ≈ 20nm, for the transition force ($F$, pN) vs change in DNA extension ($\Delta x$,nm) plots.

** Standard deviation for the $\Delta L_U$ and $\Delta L_W$ distribution, $\sigma_{\Delta L_U}$ and $\sigma_{\Delta L_W}$, respectively.

*** These experiments were performed in context 1.

’ P-value of t-student test, comparing $\Delta L_U$ and $\Delta L_W$ distribution of λ*p*$_R$ wt at 40mM KCl with the rest of conditions tested.

[++] n and N are the total number of transitions and open complexes per condition tested, respectively.

**Table S5:** Summary of the thermodynamic analysis of transitions

| | | Disruption of promoter-RNAP interactions ($u \rightarrow w$) | | | Formation of promoter-RNAP interactions ($u \rightarrow w$) | | | Change in free energy of promoter-RNAP formation | | |
|---|---|---|---|---|---|---|---|---|---|---|
| **Promoter** | **Condition** | $\langle W_{w\to u}\rangle$* | $\sigma^{W}_{w\to u}$◊ | $\langle W^{dis}_{w\to u}\rangle$+ | $\langle W_{u\to w}\rangle$* | $\sigma^{W}_{u\to w}$◊ | $\langle W^{dis}_{u\to w}\rangle$++ | $\Delta G_{u\to w}$** | $W_{strech}$ | $\Delta G^{0}_{u\to w}$*** |
| **λ*p*R wt (context 1)** | 40mM KCl | 18.7 ± 0.1 | 5.0 | 0.4 ± 0.3 | 17.9 ± 0.2 | 4.7 | 0.4 ± 0.3 | -18.3 ± 0.3 | 5.4 | -12.9 ± 0.3 |
| **λ*p*R wt (context 2)** | | 20.2 ± 0.2 | 5.0 | 0.6 ± 0.3 | 18.9 ± 0.2 | 4.4 | 0.6 ± 0.3 | -19.5 ± 0.3 | 5.8 | -13.7 ± 0.3 |
| **λ*p*R wt (Ligation)** | | 21.3 ± 0.5 | 6.4 | 0.9 ± 0.3 | 19.6 ± 0.4 | 5.4 | 0.9 ± 0.3 | -20.5 ± 0.3 | 7.1 | -13.4 ± 0.3 |
| **λ*p*R mutA****** | | 27.3 ± 0.2 | 4.2 | 0.7 ± 0.3 | 25.9 ± 0.2 | 3.8 | 0.7 ± 0.3 | -26.6 ± 0.2 | 6.3 | -20.2 ± 0.2 |
| **λ*p*R mutB****** | | 18.6 ± 0.6 | 5.6 | 2.5 ± 0.3 | 13.4 ± 0.2 | 3.2 | 2.7 ± 0.3 | -16.1 ± 0.5 | 5.7 | -10.3 ± 0.5 |
| **λ*p*R wt****** | 100mM KGlu | 24.6 ± 0.4 | 5.7 | 0.8 ± 0.5 | 23.0 ± 0.3 | 6.2 | 0.8 ± 0.4 | -23.8 ± 0.3 | 6.6 | -17.2 ± 0.3 |
| | 300mM KCl | 18.1 ± 0.2 | 3.9 | 0.6 ± 0.3 | 16.9 ± 0.2 | 4.1 | 0.6 ± 0.3 | -17.5 ± 0.2 | 6.9 | -10.6 ± 0.2 |
| | 100μM ppGpp | 16.6 ± 0.3 | 4.4 | 0.3 ± 0.4 | 16.0 ± 0.2 | 4.1 | 0.3 ± 0.4 | -16.2 ± 0.3 | 6.5 | -9.7 ± 0.3 |

* Mean irreversible work of transitions, $\langle W_{w\to u}\rangle$ (± error) and $\langle W_{u\to w}\rangle$ (± error), respectively.
◊ Standard deviation from of transition work distributions $\sigma^{W}_{w\to u}$ and $\sigma^{W}_{u\to w}$, respectively.
** Reversible work or free energy in equilibrium for transitions, $\Delta G_{u\to w}$ (± error). It was estimated using the Crooks fluctuation theorem.
+ Mean dissipated work of transitions, $\langle W^{dis}_{w\to u}\rangle = \langle W_{w\to u}\rangle - \Delta G_{w\to u}$ and $\langle W^{dis}_{u\to w}\rangle = \langle W_{u\to w}\rangle - \Delta G_{w\to u}$ (± error), respectively.
*** Reversible work at cero force, $\Delta G^{0}_{w\to u} = \Delta G_{w\to u} - W_{strech}$ (± error). $W_{strech}$ is the entropy loss due to the stretching of the DNA context and the extended λ*p*R promoter.
**** These experiments were performed in context 1.

**Table S6**: summary of transition force during the pulling/relaxing experiments per type of transition

| | | | **Disruption of promoter-RNAP interactions ($u \to w$)** | | | **Formation of promoter-RNAP interactions ($u \to w$)** | | | | |
|---|---|---|---|---|---|---|---|---|---|---|
| **Promoter** | **Condition** | **type** | $\langle F_U \rangle$* | $\sigma_{F,U}{}^{+}$ | $R^2$(U)$^{++}$ | $\langle F_W \rangle$* | $\sigma_{F,W}{}^{+}$ | $R^2(W)^{++}$ | **n**◊ | **N**◊◊ |
| **λ$p_R$ wt (Context 1)** | 40mM KCl | **Low-force** | 8.0 ± 0.1 | 0.5 | 0.94 | 7.1 ± 0.1 | 0.6 | 0.92 | 419 | 29 |
| | | **High-force** | 8.8 ± 0.1 | 0.4 | 0.93 | 8.7 ± 0.1 | 0.3 | 0.93 | 405 | |
| | | **Single** | 9.3 ± 0.1 | 0.6 | 0.96 | 9.0 ± 0.1 | 0.7 | 0.98 | 515 | |
| **λ$p_R$ wt (Context 2)** | | **High-force** | 9.9 ± 0.1 | 0.3 | 0.99 | 9.7 ± 0.1 | 0.3 | 0.98 | 178 | 15 |
| | | **Single** | 9.8 ± 0.2 | 1.1 | 0.94 | 9.3 ± 0.1 | 1 | 0.96 | 500 | |
| **λ$p_R$ wt ligation (Context 1)** | | **Single** | 9.6 ± 0.3 | 1.5 | 0.89 | 9.3 ± 0.1 | 1.0 | 0.97 | 500 | 14 |
| **λ$p_R$ mutA**** | | **Low-force** | 9.9 ± 0.2 | 1.1 | 0.84 | 8.1 ± 0.3 | 1.3 | 0.87 | 124 | 14 |
| | | **High-force** | 10.9 ± 0.1 | 0.9 | 0.98 | 10.6 ± 0.1 | 0.7 | 0.90 | 125 | |
| | | **Single** | 11.8 ± 0.2 | 1.1 | 0.86 | 11.3 ± 0.2 | 1.1 | 0.86 | 246 | |
| **λ$p_R$ mutB**** | | **Single** | 10.9 ± 0.1 | 1.2 | 0.98 | 9.0 ± 0.1 | 1.1 | 0.94 | 226 | 6 |
| **λ$p_R$ wt**** | 100mM KGlu | **Low-force** | 10.1 ± 0.1 | 0.6 | 0.97 | 8.6 ± 0.6 | 1 | 0.95 | 376 | 25 |
| | | **High-force** | 11.2 ± 0.1 | 0.5 | 0.98 | 10.9 ± 0.1 | 0.5 | 0.99 | 370 | |
| | | **Single** | 10.6 ± 0.1 | 1.1 | 0.96 | 10.2 ± 0.1 | 0.9 | 0.91 | 812 | |
| | 300mM KCl | **Low-force** | 8.6 ± 0.2 | 0.8 | 0.92 | 7.8 ± 0.1 | 1 | 0.96 | 218 | 15 |
| | | **High-force** | 10.1 ± 0.1 | 0.8 | 0.91 | 10.0 ± 0.2 | 0.8 | 0.93 | 217 | |
| | | **Single** | 9.9 ± 0.1 | 0.5 | 0.99 | 9.7 ± 0.6 | 0.5 | 0.97 | 293 | |
| | 100µM ppGpp | **Low-force** | 8.8 ± 0.1 | 0.8 | 0.97 | 7.8 ± 0.1 | 0.9 | 0.98 | 467 | 12 |
| | | **High-force** | 10.1 ± 0.2 | 0.7 | 0.84 | 10.0 ± 0.2 | 0.7 | 0.91 | 454 | |
| | | **Single** | 12.0 ± 0.2 | 0.6 | 0.88 | 11.7 ± 0.7 | 1.3 | 0.68 | 143 | |
| | | | 10.1 ± 0.4 | 0.7 | | 9.8 ± 0.3 | 0.5 | | | |

* Mean of transition forces (pN), $\langle F_U \rangle$ (± error) and $\langle F_W \rangle$ (± error) respectively.
** These experiments were performed in context 1.
+ Standard deviation of transition force (pN) distribution, $\sigma_{F,U}$ and $\sigma_{F,W}$
++ R-sq from normal fitting to the transition force (pN) distribution.
◊ Number of transitions per condition and type of transition.
◊◊ Number of open complexes.

**Table S7:** Summary of change in DNA extension transitions during the pulling/relaxing experiments per type of transition

| | | | Disruption of promoter-RNAP interactions ($u \rightarrow w$) | | | Formation of promoter-RNAP interactions ($u \rightarrow w$) | | | | |
|---|---|---|---|---|---|---|---|---|---|---|
| **Promoter** | **Condition** | **type** | $\langle \Delta x \rangle_U$* | $\sigma_{\Delta x,U}$⁺ | $R^2(U)$⁺⁺ | $\langle \Delta x \rangle_W$ | $\sigma_{\Delta x,W}$⁺ | $R^2(W)$⁺⁺ | n◊ | N◊◊ |
| **λ*p*R wt (Context 1)** | 50mM KCl | **Low-force** | 20.7 ± 0.1 | 6 | 0.94 | 18.4 ± 0.2 | 4.9 | 0.99 | 419 | 29 |
| | | **High-force** | 15.3 ± 0.4 | 4.7 | 0.95 | 15.0 ± 0.5 | 4.9 | 0.91 | 405 | |
| | | **Single** | 15.5 ± 0.2 | 3.2 | 0.98 | 15.1 ± 0.1 | 3.2 | 0.99 | 920 | |
| **λ*p*R wt (Context 2)** | | **High-force** | 13.9 ± 0.2 | 2.9 | 0.99 | 13.8 ± 0.4 | 2.7 | 0.95 | 178 | 15 |
| | | **Single** | 15.4 ± 0.4 | 2.9 | 0.94 | 15.1 ± 0.3 | 2.5 | 0.96 | 500 | |
| **λ*p*R wt ligation (Context 1)** | | **Single** | 15.7 ± 0.2 | 2.7 | 0.98 | 15.0 ± 0.1 | 2.5 | 0.99 | 500 | 14 |
| **λ*p*R mutA**** | | **Low-force** | 23.2 ± 0.2 | 2.8 | 0.98 | 17.7 ± 0.5 | 3.4 | 0.91 | 124 | 14 |
| | | **High-force** | 17.3 ± 0.5 | 2.9 | 0.9 | 17.1 ± 0.3 | 2.1 | 0.94 | 125 | |
| | | **Single** | 17.5 ± 0.2 | 1.9 | 0.96 | 17.2 ± 0.1 | 1.7 | 0.95 | 246 | |
| **λ*p*R mutB**** | | **Single** | 12.5 ± 0.8 | 3.8 | 0.89 | 11.1 ± 0.2 | 2.8 | 0.99 | 226 | 6 |
| **λ*p*R wt**** | 100mM KGlu | **Low-force** | 21.8 ± 0.3 | 3.3 | 0.96 | 19.9 ± 0.4 | 3.9 | 0.96 | 376 | 25 |
| | | **High-force** | 17.3 ± 0.2 | 3.2 | 0.99 | 17.1 ± 0.3 | 3.0 | 0.96 | 370 | |
| | | **Single** | 16.6 ± 0.4 | 3.6 | 0.95 | 16.2 ± 0.4 | 3.5 | 0.96 | 812 | |
| | 300mM KCl | **Low-force** | 19.2 ± 0.5 | 3.5 | 0.96 | 17.1 ± 0.5 | 3.0 | 0.95 | 218 | 15 |
| | | **High-force** | 13.5 ± 0.2 | 2.1 | 0.98 | 13.4 ± 0.8 | 2.8 | 0.92 | 217 | |
| | | **Single** | 13.0 ± 0.3 | 2.6 | 0.96 | 12.5 ± 0.2 | 2.4 | 0.98 | 293 | |
| | 100uM ppGpp | **Low-force** | 19.4 ± 0.1 | 3.5 | 0.99 | 15.2 ± 0.1 | 3.1 | 0.99 | 467 | 12 |
| | | **High-force** | 11.9 ± 0.1 | 2.4 | 0.99 | 11.8 ± 0.2 | 2.6 | 0.97 | 454 | |
| | | **Single** | 12.1 ± 0.2 | 2.9 | 0.98 | 11.3 ± 0.2 | 2.5 | 0.97 | 143 | |

* Mean of change in DNA extension of transitions (nm), $\langle \Delta x \rangle_U$ (± error) and $\langle \Delta x \rangle_W$ (± error), respectively.
** These experiments were performed in context 1.
⁺ Standard deviation of the change in DNA extension transition (nm) distribution, $\sigma_{\Delta x,U}$ and $\sigma_{\Delta x,W}$
⁺⁺ $R^2$ from normal fitting to change in DNA extension transition (nm) distribution, $R^2(U)$ and $R^2(W)$, respectly.
◊ Number of transitions and type of transition.
◊◊ Number of open complexes.

**Table S8:** EM data collection and SPA

| | **EMD-0340** |
|---|---|
| Magnification | 60,000 |
| Voltage (kV) | 200 |
| Electron square ($e^-$/Å$^2$) | 20 |
| Defocus range (µm) | -1 to -2 |
| Pixel size (Å) | 1.78 |
| Symmetry imposed | C1 |
| Initial particles images | 60,393 |
| Final particles images | 16,015 |
| Map resolution (Å) | 17 |
| FSC-Threshold | ½-bit |

**Table S9:** List of oligonucleotides to form 100-λ$p_R$-Cy3, mutA-Cy3 and mutB-Cy3 and 40-λ$p_R$-Cy3 constructs

| **Oligonucleotide** | **sequence** |
|---|---|
| downstream-sense$^{Cy3}$ | 5-/phos/GATAATGGTTGC<u>A</u>TGTACTAAGGAGGTTGT-3 |
| downstream-antisense | 5-ACAACCTCCTTAGTACATGCAACCAT-3 |
| upstream-sense | 5-GCGTGTTGACTATTTTACCTCTGGCGGT-3 |
| upstream-antisense | 5-/phos/TATCACCGCCAGAGGTAAAATAGTCAACACGC-3 |
| startUP-sense | 5-/phos/GGGATAAATATCTAACACCGTGCGTGTTGACTATTTTACCTCTGGCGGT-3 |
| startUP-antisense | 5-/phos/TATCACCGCCAGAGGTAAAATAGTCAACACGCACGGTGTTAGATATTTA-3 |
| endUP-sense | 5-TTTCTTTTTTGTGCTCATACGTTAAATCTATCACCGCAA-3 |
| endUP-antisense | 5-/phos/TCCCTTGCGGTGATAGATTTAACGTATGAGCACAAAAAAGAAA-3 |
| startUP-sense | 5-/phos/GGGATAAATATCTAACACCGTGCGTGTTGACTATTTTACCTCTGGCGGT-3 |
| startUP-antisense | 5-/phos/TATCACCGCCAGAGGTAAAATAGTCAACACGCACGGTGTTAGATATTTA-3 |
| mutA-sense | 5-ATGATTACGCCAAGCTTGCATTTAAATCTATCACCGCAA-3 |
| mutA-antisense | 5-/phos/TCCCTTGCGGTGATAGATTTAAATGCAAGCTTGGCGTAATCAT-3 |
| mutB$^{I}$-sense | 5-/phos/CCTGGGGTGCCTAGCGTGTGATCTATTGACTATTTTACCTCTGGCGGT-3 |
| mutB$^{I}$-antisense | 5-/phos/TATCACCGCCAGAGGTAAAATAGTCAATAGATCACACGCTAGGCACCC-3 |
| mutB$^{II}$-sense | 5-ATGATTACGCCAAGCTTGCATGCCTGCAGGTTTAAACAGT-3 |
| mutB$^{II}$-antisense | 5-/phos/CAGGACTGTTTAAACCTGCAGGCATGCAAGCTTGGCGTAATCAT-3 |

**Table S10:** Individual kinetic constants ($s^{-1}$) of RNAP-λ$p_R$ association ($k_a$), DNA bubble formation ($k_o$), and promoter escape ($k_e$); and their fitting parameters per repetition.

| Construct | Condition | $k_a$ ($s^{-1}$)* | $k_o$ ($s^{-1}$)* | RMSE | $R^2$ | $k_e$* | RMSE | $R^2$ |
|---|---|---|---|---|---|---|---|---|
| **40-λ$p_R$-Cy3** | **low salt** | $8.06 \times 10^{-2}$ | $3.74 \times 10^{-4}$ | 0.0099 | 0.99 | $1.74 \times 10^{-2}$ | 0.0111 | 0.69 |
| | | $1.31 \times 10^{-2}$ | $3.84 \times 10^{-4}$ | 0.0096 | 0.99 | $5.33 \times 10^{-2}$ | 0.0111 | 0.78 |
| | | $6.86 \times 10^{-2}$ | $2.64 \times 10^{-4}$ | 0.0097 | 0.99 | $4.05 \times 10^{-2}$ | 0.0094 | 0.79 |
| | | $1.34 \times 10^{-2}$ | $3.18 \times 10^{-4}$ | 0.0097 | 0.98 | ND | ND | ND |
| | | $1.18 \times 10^{-2}$ | $5.87 \times 10^{-4}$ | 0.0091 | 0.99 | $2.70 \times 10^{-2}$ | 0.0094 | 0.57 |
| | **ppGpp** | $0.67 \times 10^{-2}$ | $0.38 \times 10^{-4}$ | 0.0092 | 0.99 | $0.9 \times 10^{-3}$ | 0.0104 | 0.83 |
| | | $1.05 \times 10^{-2}$ | $1.04 \times 10^{-4}$ | 0.0103 | 0.99 | $2.59 \times 10^{-2}$ | 0.0117 | 0.86 |
| | | $1.01 \times 10^{-2}$ | $0.17 \times 10^{-4}$ | 0.0112 | 0.99 | $2.45 \times 10^{-2}$ | 0.0111 | 0.88 |
| | | ND | ND | ND | ND | $1.85 \times 10^{-2}$ | 0.0111 | 0.84 |
| **100-λ$p_R$-Cy3** | **low salt** | $1.78 \times 10^{-2}$ | $2.47 \times 10^{-3}$ | 0.0136 | 0.99 | $1.24 \times 10^{-2}$ | 0.0152 | 0.86 |
| | | $1.65 \times 10^{-2}$ | $3.52 \times 10^{-3}$ | 0.0144 | 0.99 | $2.13 \times 10^{-2}$ | 0.0152 | 0.92 |
| | | $1.51 \times 10^{-2}$ | $2.28 \times 10^{-3}$ | 0.0147 | 0.99 | $1.97 \times 10^{-2}$ | 0.0158 | 0.78 |
| | | $1.60 \times 10^{-2}$ | $1.81 \times 10^{-3}$ | 0.0136 | 0.99 | $2.42 \times 10^{-2}$ | 0.0170 | 0.77 |
| | **high salt** | 2.48 | $5.98 \times 10^{-4}$ | 0.0137 | 0.99 | $1.44 \times 10^{-2}$ | 0.0144 | 0.93 |
| | | 3.13 | $6.54 \times 10^{-4}$ | 0.0132 | 0.99 | $1.22 \times 10^{-2}$ | 0.0144 | 0.96 |
| | | 5.19 | $6.26 \times 10^{-4}$ | 0.0205 | 0.99 | $1.21 \times 10^{-2}$ | 0.0152 | 0.95 |
| | | 2.20 | $9.97 \times 10^{-4}$ | 0.0138 | 0.99 | $1.46 \times 10^{-2}$ | 0.0162 | 0.95 |
| | **glutamate** | $0.88 \times 10^{-2}$ | $2.07 \times 10^{-3}$ | 0.0139 | 0.99 | $2.13 \times 10^{-2}$ | 0.0132 | 0.93 |
| | | $1.07 \times 10^{-2}$ | $3.32 \times 10^{-3}$ | 0.0122 | 0.99 | $2.43 \times 10^{-2}$ | 0.0133 | 0.93 |
| | | $1.02 \times 10^{-2}$ | $2.62 \times 10^{-3}$ | 0.0129 | 0.99 | $1.98 \times 10^{-2}$ | 0.0155 | 0.92 |
| | | ND | ND | ND | ND | $3.16 \times 10^{-2}$ | 0.0155 | 0.88 |
| | **ppGpp** | $1.82 \times 10^{-2}$ | $1.34 \times 10^{-3}$ | 0.0219 | 0.99 | $2.22 \times 10^{-2}$ | 0.0178 | 0.86 |
| | | $1.58 \times 10^{-2}$ | $1.17 \times 10^{-3}$ | 0.0161 | 0.99 | $1.53 \times 10^{-2}$ | 0.0176 | 0.88 |
| | | $1.44 \times 10^{-2}$ | $0.65 \times 10^{-3}$ | 0.0162 | 0.99 | $0.26 \times 10^{-2}$ | 0.0163 | 0.94 |
| | | $2.11 \times 10^{-2}$ | $1.08 \times 10^{-3}$ | 0.0180 | 0.99 | ND | ND | ND |
| **mutA-Cy3** | **Low salt** | $3.00 \times 10^{-2}$ | $2.64 \times 10^{-3}$ | 0.0144 | 0.99 | $1.88 \times 10^{-2}$ | 0.0139 | 0.90 |
| | | $4.20 \times 10^{-2}$ | $6.38 \times 10^{-3}$ | 0.0226 | 0.99 | $2.32 \times 10^{-2}$ | 0.0141 | 0.93 |
| | | $4.27 \times 10^{-2}$ | $4.77 \times 10^{-3}$ | 0.0218 | 0.98 | $0.91 \times 10^{-2}$ | 0.0145 | 0.94 |
| | | $3.35 \times 10^{-2}$ | $2.07 \times 10^{-3}$ | 0.0142 | 0.99 | ND | ND | ND |
| **mutB-Cy3** | | $6.31 \times 10^{-3}$ | $1.28 \times 10^{-3}$ | 0.0122 | 0.99 | $1.1 \times 10^{-2}$ | 0.0114 | 0.95 |
| | | $4.72 \times 10^{-3}$ | $1.34 \times 10^{-3}$ | 0.0151 | 0.99 | $1.3 \times 10^{-2}$ | 0.0132 | 0.94 |
| | | $7.37 \times 10^{-3}$ | $1.76 \times 10^{-3}$ | 0.0160 | 0.99 | $1.6 \times 10^{-2}$ | 0.0165 | 0.97 |
| | | $6.24 \times 10^{-3}$ | $1.19 \times 10^{-3}$ | 0.0217 | 0.99 | $1.4 \times 10^{-2}$ | 0.0148 | 0.94 |
| | | $4.77 \times 10^{-3}$ | $0.69 \times 10^{-3}$ | 0.0128 | 0.99 | $1.3 \times 10^{-2}$ | 0.0132 | 0.97 |

RMSE: root mean square error.

ND: not determined.

**Table S11:** kinetic constants ($s^{-1}$) of RNAP-λ$p_R$ association ($k_a$), DNA bubble formation ($k_o$), and promoter escape ($k_e$).

| Construct | Condition | $\Delta G^0_{u\rightarrow w}$** (kcal/mol) | $k_a$ ($s^{-1}$)* | $k_o$ ($s^{-1}$)* | n+ | $k_e$* | n++ |
|---|---|---|---|---|---|---|---|
| **40-λ$p_R$-Cy3** | **low salt** | N.D | (1.1 ± 0.1)x$10^{-2}$ | (3.9 ± 0.5)x$10^{-4}$ | 5 | (3.5 ± 0.7)x$10^{-2}$ | 4 |
| | **ppGpp** | | (0.9 ± 0.1)x$10^{-2}$ | (0.5 ± 0.2)x$10^{-4}$ | 3 | (2.0 ± 0.2)x$10^{-2}$ | 4 |
| **100-λ$p_R$-Cy3** | **low salt** | -12.9 ± 0.3 | (1.6 ± 0.1)x$10^{-2}$ | (25.2 ± 3.6)x$10^{-4}$ | 4 | (1.9 ± 0.3)x$10^{-2}$ | 4 |
| | **high salt** | -10.6 ± 0.2 | 32.5 ± 6.8 | (7.2 ± 0.9)x$10^{-4}$ | 4 | (1.3 ± 0.1)x$10^{-2}$ | 4 |
| | **glutamate** | -17.2 ± 0.3 | (1.0 ± 0.1)x$10^{-2}$ | (26.7 ± 3.6)x$10^{-4}$ | 3 | (2.4 ± 0.3)x$10^{-2}$ | 4 |
| | **ppGpp** | -9.7 ± 0.3 | (1.7 ± 0.1)x$10^{-2}$ | (10.6 ± 1.5)x$10^{-4}$ | 4 | (1.3 ± 0.6)x$10^{-2}$ | 3 |
| **mutA-Cy3** | **Low salt** | -20.2 ± 0.2 | (3.7 ± 0.3)x$10^{-2}$ | (39.7 ± 9.9)x$10^{-4}$ | 4 | (1.7 ± 0.4)x$10^{-2}$ | 3 |
| **mutB-Cy3** | | -10.3 ± 0.5 | (0.6 ± 0.1)x$10^{-2}$ | (12.5 ± 1.7)x$10^{-4}$ | 5 | (0.9 ± 0.4)x$10^{-2}$ | 5 |

* $k_b$, $k_o$ and $k_e$ are the RNAP-λ$p_R$ association ($s^{-1}$), DNA bubble formation ($s^{-1}$) and promoter escape ($s^{-1}$) rate constants, respectively. Values are reported as mean ± SEM (standard error of the mean)

** Energy associated to upstream and downstream processes in the open complex, $\Delta G^0_{u\rightarrow w}$ (supplementary Table 5).

+ Number of trajectories of open complex formation considered in the analysis to extract $k_a$ and $k_o$.

++ Number of trajectories of promoter escape considered in the analysis to extract $k_e$.
